# Hippocampus has lower oxygenation and weaker control of brain blood flow than cortex, due to microvascular differences

**DOI:** 10.1101/835728

**Authors:** K. Shaw, L. Bell, K. Boyd, D.M. Grijseels, D. Clarke, O. Bonnar, H.S. Crombag, C.N. Hall

## Abstract

The hippocampus is essential for spatial and episodic memory but is damaged early in Alzheimer’s disease and is very sensitive to hypoxia. Understanding how it regulates its oxygen supply is therefore key for designing interventions to preserve its function. However, studies of neurovascular function in the hippocampus *in vivo* have been limited by its relative inaccessibility. Here we compared hippocampal and visual cortical neurovascular function in awake mice, using two photon imaging of individual neurons and vessels and measures of regional blood flow and haemoglobin oxygenation. We show that blood flow, blood oxygenation and neurovascular coupling were decreased in the hippocampus compared to neocortex, because of differences in both the vascular network and pericyte and endothelial cell function. Modelling oxygen diffusion indicates that these features of the hippocampal vasculature could explain its sensitivity to damage during neurological conditions, including Alzheimer’s disease, where the brain’s energy supply is decreased.

## Introduction

Brain processes that depend on the hippocampus become dysfunctional early in several disease states, including Alzheimer’s disease, as well as during ageing and after lifestyle changes such as the adoption of a high fat diet ^1–3^. Changes in vascular function are also features of early stages of these conditions ^2^. CA1 pyramidal cells of the hippocampus are particularly susceptible to death after a hypoxic insult ^4^, suggesting that maintaining an adequate oxygen supply may be particularly vital for this brain region.

Physiologically, the brain finely regulates its oxygen supply by a process called “neurovascular coupling” whereby active neurons signal to dilate local blood vessels, increasing blood flow and the supply of oxygen and glucose to these active brain regions. In the neocortex, neurovascular coupling produces an increase in blood flow that more than compensates for the increase in oxygen consumed by active neurons, generating an overall increase in blood oxygenation in the active brain region. The associated decrease in levels of deoxygenated haemoglobin underlies the positive blood oxygen level-dependent (BOLD) signal. In the neocortex, this increase in BOLD correlates with neuronal activity, enabling it to be used as a surrogate measure of brain activation in functional magnetic resonance imaging (fMRI) studies. In the hippocampus, however, the BOLD signal does not seem to reliably correlate with neuronal activity ^5^, as local field potential changes occur without measurable BOLD signals ^6^. This not only suggests that fMRI studies measuring hippocampal BOLD are less sensitive than cortex to increases in neuronal activity, but also that the hippocampus may be physiologically less able to increase its oxygen supply, and this may underlie its sensitivity to damage at the onset of pathophysiological conditions.

However, because the hippocampus lies beneath the neocortex, and is therefore less accessible, previous studies have not been able to directly measure neurovascular coupling in this region *in vivo*, and therefore no direct evidence exists whether the hippocampus is indeed less able to regulate its energy supply. By removing overlying cortex ^7^, we could implant a cranial window over the hippocampus and record neuronal and vascular activity using two-photon imaging, laser doppler flowmetry and haemoglobin spectroscopy to compare neurovascular coupling in dorsal CA1 to that in the visual cortex. We found that the hippocampus had lower resting blood flow and blood oxygenation compared to visual cortex, which was due both to a lower capillary density and reduced red blood cell velocity in individual capillaries than in visual cortex.

We then studied how well blood vessels could respond to increases in local excitatory neuronal activity and found that not only did blood vessels in the hippocampus dilate less frequently to local increases in activity than in visual cortex, but when responses did occur, the dilations were smaller. Pericyte morphology and vascular expression of proteins that mediate dilation were significantly different in the hippocampal capillary bed compared to cortex, suggesting that the hippocampal vasculature is less able to dilate to similar increases in neuronal activity. To understand how these vascular differences impact neuronal oxygen availability, we modelled oxygen diffusion from vessels. Our results indicated that oxygen becomes limiting for ATP synthesis in tissue furthest from blood vessels in the hippocampus, but not the visual cortex. We propose that this decreased neurovascular function in the hippocampus contributes to its vulnerability to damage in disease states.

## Results

We measured neurovascular coupling in the dorsal CA1 region of the hippocampus (HC) and in visual cortex (V1) of head-fixed mice expressing GCaMP6f in excitatory (glutamatergic) neurons ^8^. Mice could run on a running cylinder or remain stationary, while visual stimuli (drifting gratings or a virtual reality environment) were presented (Online Methods, Figures 1a-d). In some experiments, combined laser doppler flowmetry/haemoglobin spectroscopy was used to record levels of deoxy-(Hbr) and oxyhaemoglobin (HbO), and cerebral blood flow (CBF) (oxy-CBF probe; Figure 1e), allowing us to calculate regional blood oxygen saturation (SO_2_) and the cerebral metabolic rate of oxygen consumption (CMRO_2_; Figure 1f). In other experiments, individual neurons and blood vessels were imaged using two-photon microscopy (e.g. Figure 3a).

**Figure 1:**
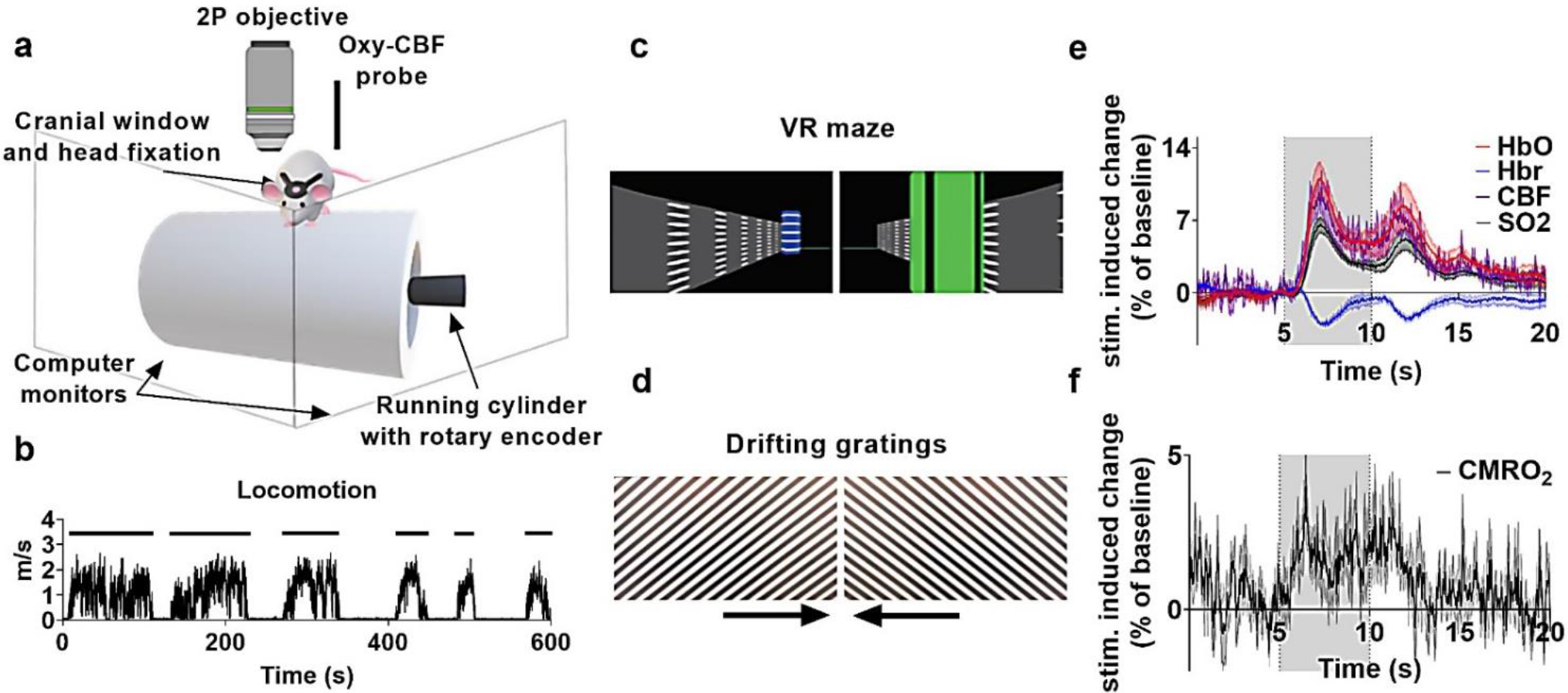
Experimental set-up and example haemodynamic data. **(a)** Schematic of the imaging set-up. Either the two-photon objective or oxy-CBF probe were used to collect data while the mouse was head-fixed but awake and able to run on the cylinder. **(b)** Representative locomotion recorded by the rotary encoder during one imaging session (metres per second). Distinct periods of running are indicated by the black bars. A virtual reality maze **(c)** or drifting gratings **(d)** were presented on the screens in (a). Locomotion advanced the mice through the virtual reality maze. The arrows beneath the drifting gratings display show the direction the gratings travelled. **(e)** Example haemodynamic recordings from visual cortex using the oxy-CBF probe during visual stimulation (grey bar represents stimulation, N=4 animals, 10 sessions, 202 trials). **(f)** The cerebral metabolic rate of oxygen consumption (CMRO_2_) is calculated from the haemodynamic parameters collected using the oxy-CBF probe for the data in (e) (see Online Methods). All data traces are unsmoothed averages, and error bars represent mean +/− SEM.

### Blood flow and oxygenation are lower at rest in hippocampus than V1

We first compared haemodynamics in HC and V1 in the absence of visual stimulation when the mouse was stationary. CMRO_2_, reflecting energy use by summed neuronal activity, was not different between regions (Figure 2a). However, despite similar energy demands, the resting CBF and SO_2_ were significantly lower in HC (Figures 2b, c). In part, these differences in net blood flow and oxygenation arise from a lower capillary density in HC than cortex (Figures 2d, e ^9^). However, when we measured red blood cell (RBC) velocity in individual blood vessels, after loading vessels with fluorescent dextran (Figure 2f), we found that despite the vessels themselves being of equal size (Figure 2g), RBC velocity was significantly lower in the HC than V1 (Figure 2h). This combination of a lower capillary density and RBC velocity in the HC explains both the observed lower net flow and, because fractionally more oxygen is extracted from capillaries with lower flow rates ^10^, the lower blood oxygenation we measured in HC.

**Figure 2:**
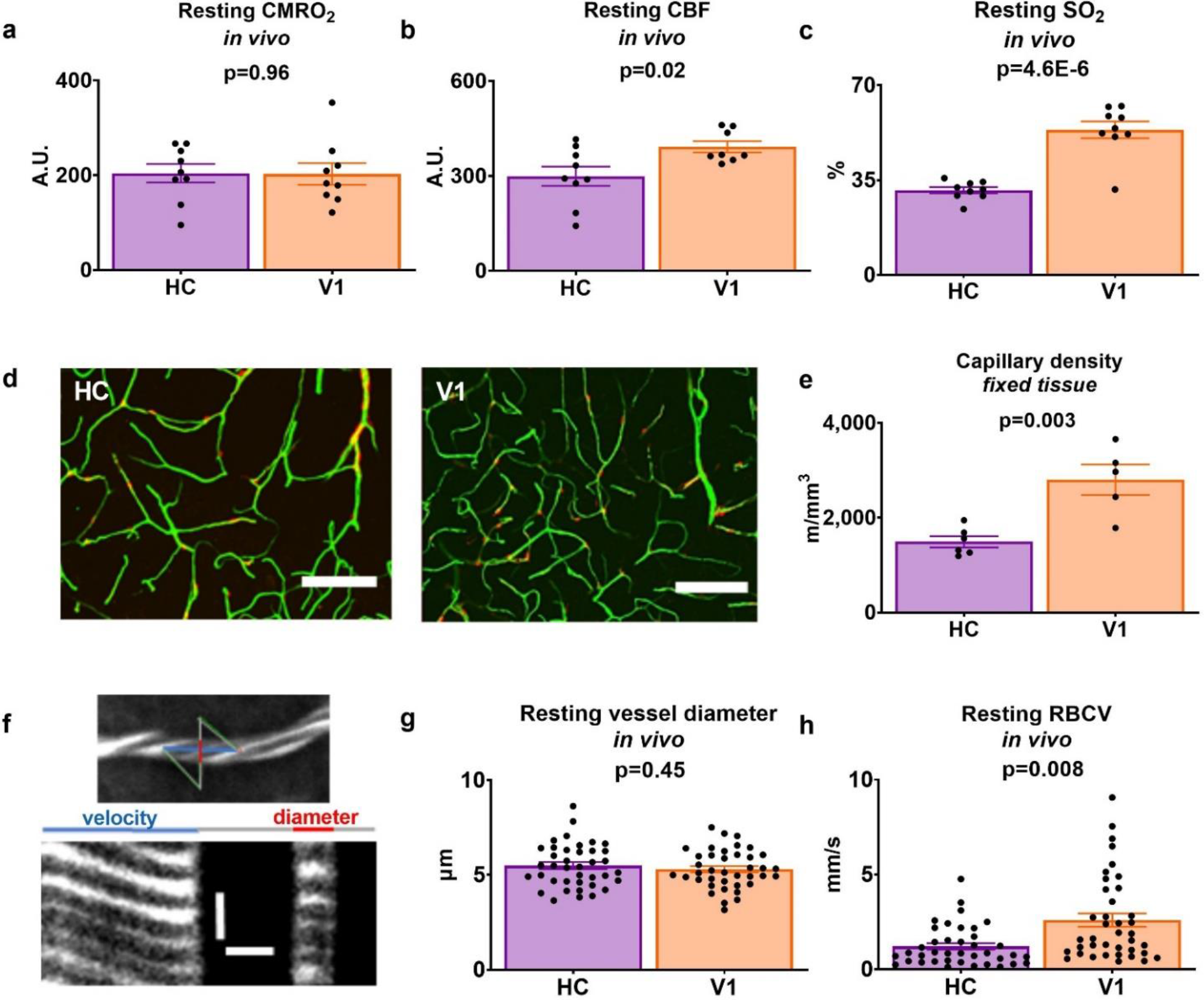
Baseline haemodynamics in HC and V1. CMRO_2_ **(a)**, CBF **(b)**, and oxygen saturation (SO_2_) **(c)** in HC (purple) and V1 (orange) in stationary mice in the dark (HC: 9 animals across 21 sessions; V1: 9 animals across 17 sessions; data points are averages for each animal). **(d)** FITC-gelatin filled vessels (green) from one NG2-DsRed mouse (pericytes are red). Scale bars represent 100 μm. Images are projections of 100 μm Z-stacks. **(e)** Capillary density in these slices was significantly lower in HC (N=6 mice) compared to V1 (N=5 mice). **(f)** Example line scan trajectory from one in vivo two-photon recording (top). The scan path of the laser goes along (blue) and across (red) the capillary (labelled with i.v. Texas Red dextran, white. Dark stripes are RBCs). Each row in the corresponding line scan image (bottom) represents a single time point. As RBCs move, their shadow shifts along the vessel, so the angle of the stripes (left, under blue line) shows how fast they are moving. The vessel diameter can be measured from the intensity profile of the Texas Red-labelled lumen (right, under red line). The vertical scale bar represents 5 ms, and the horizontal scale bar 5 μm. The diameters of these capillaries were not significantly different between regions **(g)**, but their RBC velocity at rest **(h)** was faster in V1 (39 vessels from 11 mice) than in HC (39 vessels from 9 mice). Data in bar charts represent mean +/− SEM, dots are individual vessels or mice, as indicated. P-values are from independent sample t-tests.

### Blood vessels in CA1 dilate less to local neuronal activity than those in V1

Because fMRI studies suggest neuronal activity might be less well matched to blood flow in HC than cortex, we investigated the capacity of blood vessels to dilate in response to local excitatory neuronal calcium events *in vivo* (Figures 3a-e). In both regions, vessels dilated shortly after neuronal calcium events (Figures 3d, e, g, h), however, the frequency and size of dilations were significantly greater in V1 than in HC (Figures 3c, e, h). The size of the calcium peaks was similar between regions (Figures 3d, g), suggesting that neuronal activity, and therefore energy requirements, were the same in each region during calcium events. Most of our dilations are related to calcium events and not due to random vasomotion because when traces were shuffled so that vessel dilation data was no longer aligned to calcium events, dilations occurred less frequently (V1: 5.19%, HC: 5.67% of the time; Figures 3f, i). Even when vessels did dilate in HC, the size of these responses was smaller than in V1, though the calcium events that triggered these responses were the same size (Figure 3j-m). Therefore, smaller dilations in HC were due to both a decrease in responsiveness of vessels and to dilations being smaller in HC when they did occur.

**Figure 3:**
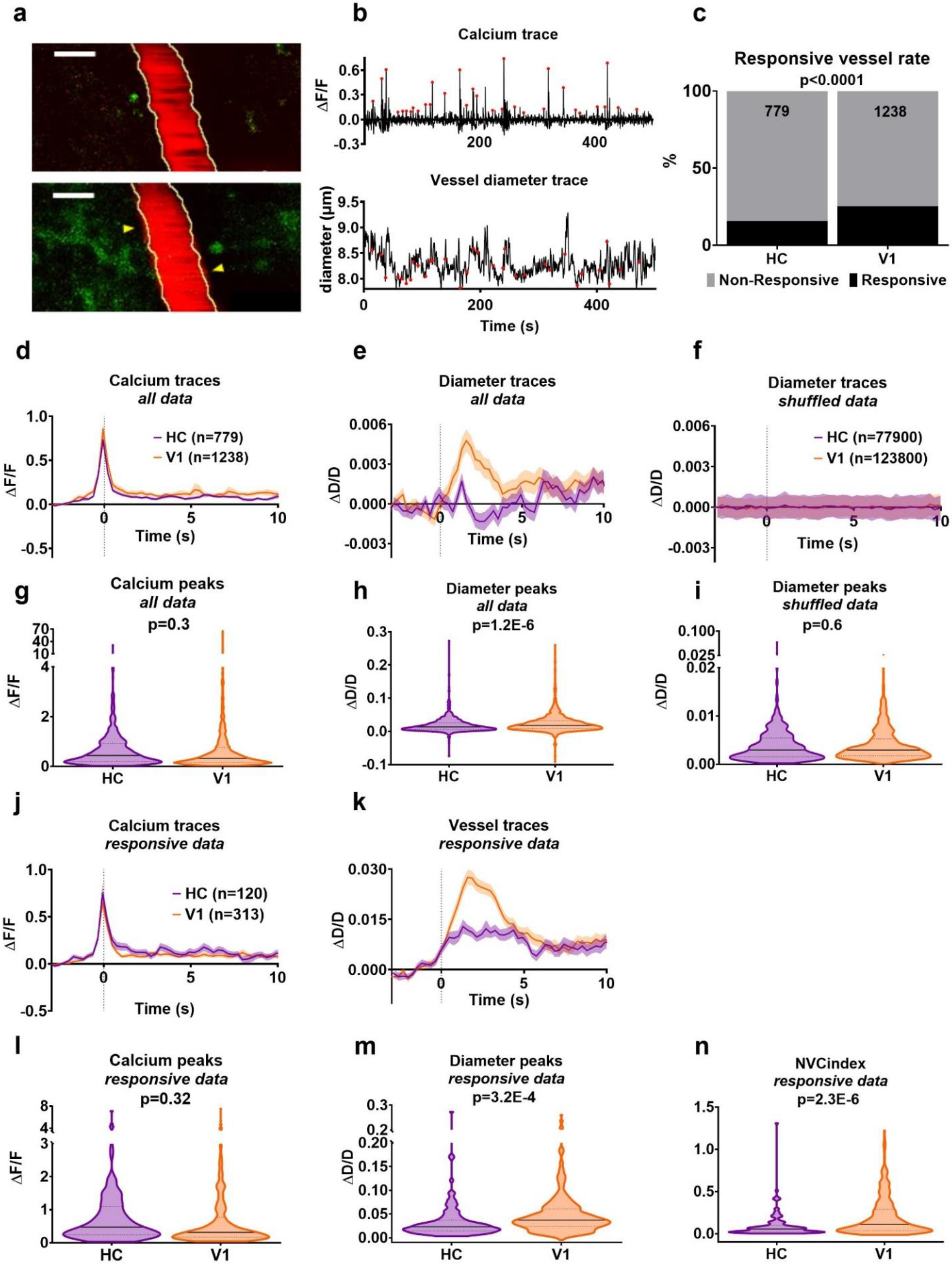
Vessel responses to local neuronal calcium events. **(a)** Texas Red dextran-filled vessel (red) and GCaMP6f-positive pyramidal neurons (green) from one recording in V1 before (top) and during (bottom) increases in neuronal calcium. Yellow outlines show vessel before calcium event. Arrows indicate largest dilation. Scale bars represent 5 μm. **(b)** Neuronal calcium averaged over all cells in the field of view of one imaging session and the corresponding vessel diameter. Red dots mark calcium events. **(c)** Vessel responses to preceding calcium events were more frequent in V1 than HC (Chi-square test). **(d)** Average calcium response and **(e)** diameter change in HC (purple, N=779 trials, 46 vessels, 6 animals) and V1 (orange, N=1238 trials, 41 vessels, 7 animals). **(f)** Diameters when vessel traces were shuffled 100 times so no longer aligned to calcium events. **(g)** There was no regional difference in the size of calcium events. **(h)** Vessel dilations were larger in V1 than HC when aligned to calcium events, but not when shuffled **(i)**. **(j)** Calcium events which led to dilations in HC (N=120 events, 6 mice) and V1 (N=313 events, 7 mice). **(k)** Corresponding diameter changes. **(l)** Calcium peaks leading to dilations were no different between regions. **(m)** Diameter peaks were significantly larger in V1 than HC. **(n)** A neurovascular coupling index (NVC_idx_) was calculated by dividing each dilation peak by its corresponding calcium peak. NVC_idx_ was lower in HC than V1. P-values are from independent sample t-tests, unless stated. Averages show mean +/− SEM. Horizontal lines on violin plots show median and interquartile range.

To characterise the relationship between each individual calcium event and corresponding vessel response, we calculated a “neurovascular coupling index” (NVC_idx_) by dividing responding vessel diameter peaks by the corresponding neuronal calcium peak, so a large NVC_idx_ represents a large dilation in response to a given change in calcium. NVC_idx_ was significantly higher in V1 than HC (Figure 3n), suggesting a decreased ability of the hippocampal vasculature to match increased blood flow to increased oxygen use compared to cortex. This reduced ability to match blood supply to changing energetic demands in the HC could be due to a decrease in the “instruction” from the neurons to tell blood vessels to dilate, or a decreased ability of HC blood vessels to respond to this signal.

### HC neurons do not express lower synthetic enzymes for vasodilatory signalling pathways

HC neurons could be less able to instruct the vasculature to dilate if HC neurons and astrocytes are less able to produce vasodilatory messengers than similar cells in V1. We tested for differences in expression levels of vasodilatory second messenger pathways in neurons and astrocytes between regions, using an open access single cell mRNA dataset from neocortex (S1) and HC (CA1) ^11^. There were no differences in levels of mRNA in individual pyramidal cells, interneurons and astrocytes of the synthetic enzymes for prostaglandins, EETs, and nitric oxide that could explain neurovascular coupling being weaker in HC than V1 (Supplementary Tables 1-3). In fact, there were significantly higher levels of Nos1 (Figure 4a) and Ptges3 (Figure 4b) expression in HC pyramidal neurons, and of Ptges3 (Figure 4c) in HC astrocytes, which would predict that, if anything, the production of vasodilatory molecules (prostaglandin and nitric oxide) would be greater in HC than V1.

**Figure 4:**
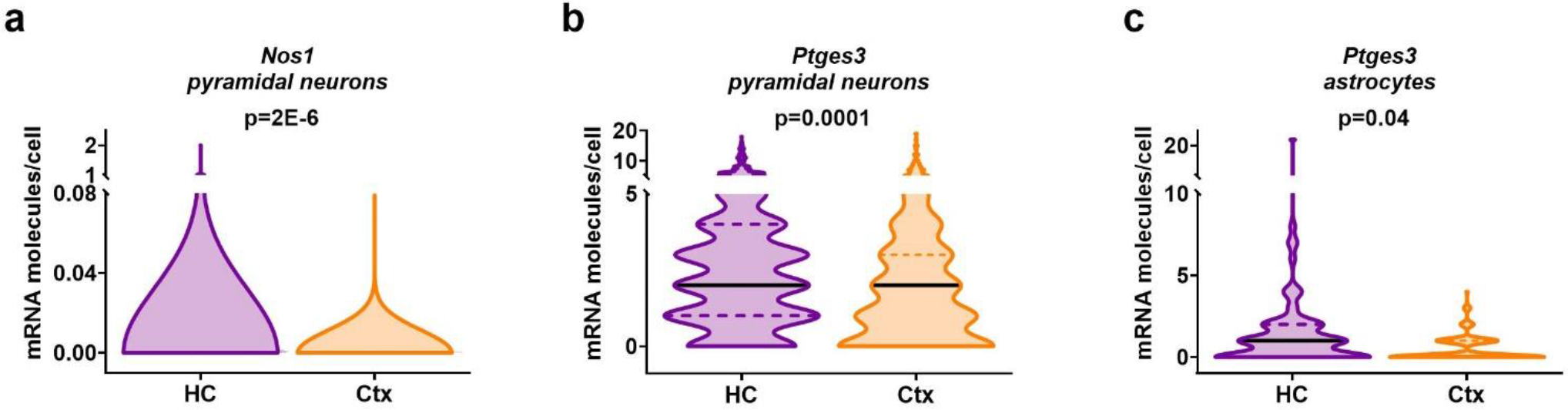
Vasodilatory second messenger pathways in HC and cortex. The expression of **(a)** Nos1 and **(b)** Ptges3 were higher in the pyramidal cells of HC (941 cells) than cortex (398 cells). **(c)** Ptges3 expression was also higher in astrocytes in HC (80 cells) than V1 (143 cells). Horizontal lines on violin plots show median and interquartile range. P-values represent the result of independent sample t-tests, with Holm-Bonferroni corrections for the multiple comparisons presented in each of Supplementary Tables 1-3.

### HC neurovascular coupling is not lower because neuronal firing is less synchronous

Alternatively, HC neurons could be less effective at signalling to blood vessels if their firing is less synchronous than in V1, so that levels of dilatory second messengers (e.g. nitric oxide or prostaglandin) summate less, so their concentration and potential effect on the vasculature is reduced. Indeed, coding in HC is sparser and distributed, while that in V1 is retinotopic, so it might be predicted that neuronal firing in HC would, indeed, be less synchronised. We tested this by imaging across a large field of view to capture large numbers of excitatory cells (Figure 5a). We tested for synchronous firing in two ways: 1) by investigating whether the cells fired at the same time, using cross-correlation of the trace from each cell with the others (Figure 5c), and 2) by measuring whether peaks in the average calcium trace across all cells (i.e. bursts of synchronous activity) were larger in V1 than in HC (Figures 5b, d). There were no significant differences in either the correlation of firing, or the average size of population calcium peaks between the regions.

**Figure 5:**
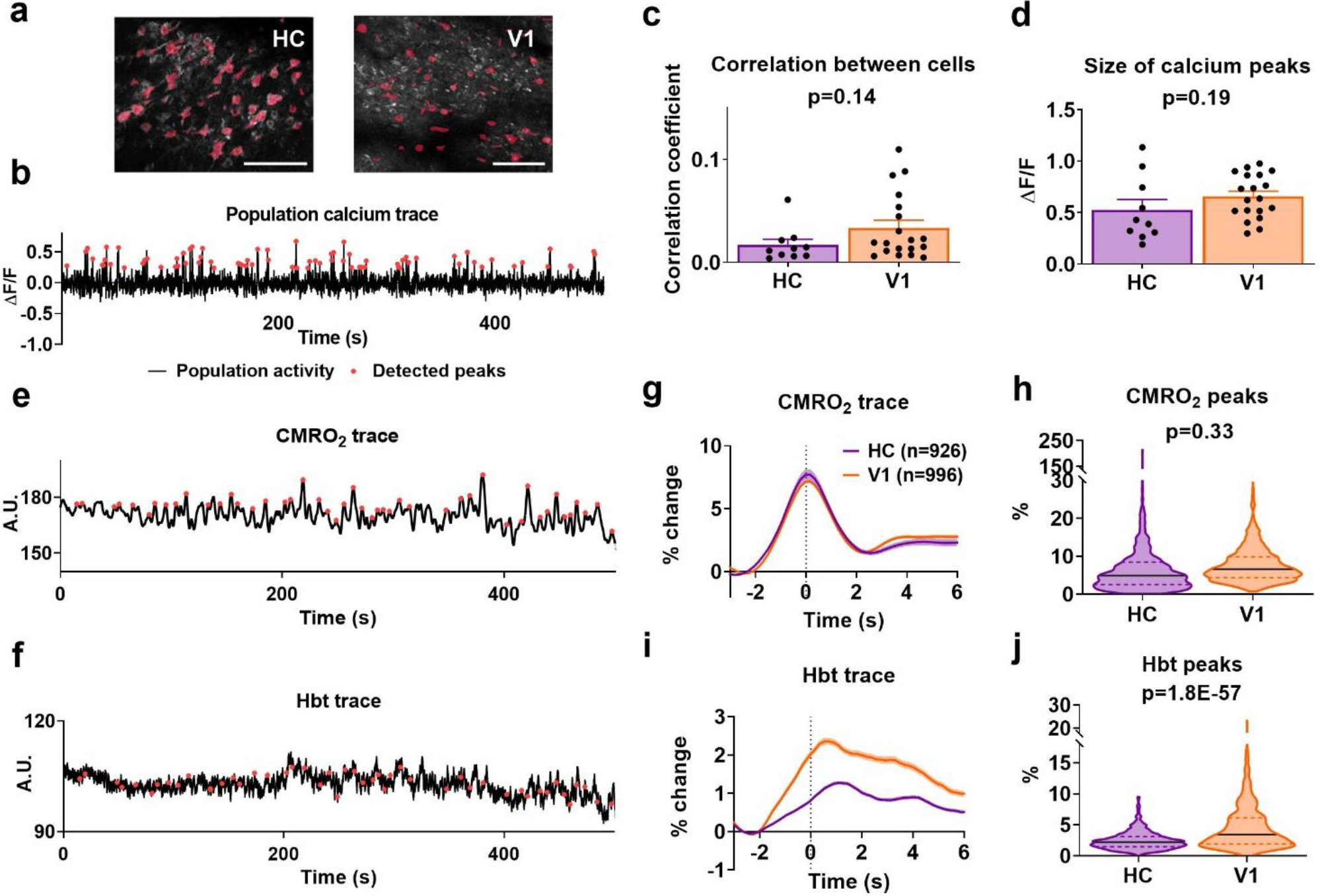
Wide-field neuronal activity patterns. **(a)** Representative wide field of view recording of calcium signals (white; maximum projection across time) in HC (left) and V1 (right). Regions of interest (ROIs) used to measure cell activity are displayed in pink. Scale bar represents 100 μm. **(b)** Example neuronal calcium trace, averaged across all detected ROIs. Times when net activity peaks (>2 SD above baseline mean) are shown by red dots. **(c)** The correlation (Pearson’s R) between individual cells within a field of view was not different between the HC (10 recordings from 5 animals, 559 cells detected in total) and V1 (19 recordings from 3 animals, 338 cells in total). **(d)** The size of net calcium peaks was also not different between regions. Dots in (c) and (d) represent separate recording sessions. **(e)** Example CMRO_2_ fluctuations over time and **(f)** corresponding total haemoglobin (Hbt). CMRO_2_ peaks are marked in red. **(g)** Averaged CMRO_2_ peaks in HC (data represents 926 peaks, taken from 9 animals across 37 recordings) and V1 (996 peaks taken from 10 animals across 46 recordings). **(h)** There was no difference in the magnitude of CMRO_2_ peaks between regions. **(i)** Hbt in response to CMRO_2_ peaks in HC and V1. **(j)** Hbt increased more in V1 than in HC within 5 s of an increase in CMRO_2_. P-values represent the result of independent sample t-tests. Bar charts and traces show mean +/− SEM, violin plots show median and interquartile range.

An alternative measure of net neuronal activity is provided by the CMRO_2_ signal. We detected peaks in CMRO_2_ (Figure 5e), alongside corresponding regional increases in cerebral blood volume due to vascular dilation (reflected in the total blood volume, Hbt; Figure 5f). In accordance with an equivalent degree of synchronous firing in V1 and HC, the sizes of the peaks in the CMRO_2_ signal were not different between regions (Figures 5g, h), yet associated cerebral blood volume changes were smaller in HC than V1 (Figures 5i, j), consistent with weaker neurovascular coupling in HC.

We wondered whether the cellular or laminar organisation of HC and V1 could explain the different neurovascular coupling properties, perhaps if second messengers released from different neuronal compartments have differential effects on the vasculature. To this end, we tested whether neurovascular coupling was different in vessels in response to calcium signals in the neuropil or nearby somas (Supplementary Figure 1). We found that, while vessels in V1 were more likely to respond to calcium signals from the neuropil, vessel responsiveness in HC did not distinguish between the different neuronal compartments. The size of the calcium peaks from neuropil or soma were no different in either HC or V1, nor were corresponding vessel responses to these calcium signals, resulting in a comparable NVC_idx_ between the different neuronal compartments within both regions, and a significantly greater NVC_idx_ in V1 than HC for both neuropil and soma. We also investigated whether there were laminar differences in neurovascular coupling that could explain our results. We found some differences in responses between layers in both regions (Supplementary Figure 2), but NVC_idx_ was significantly greater in both V1 layers than in HC.

### Pericytes have a less contractile morphology in HC than V1

In the absence of clear differences in neuronal firing properties or expression of neurovascular coupling signalling molecules, we next investigated whether our results could reflect differences in vascular function. The morphology of vascular mural cells (smooth muscle cells and pericytes) is linked to their ability to constrict and dilate blood vessels ^12^. Broadly, more contractile cells are shorter in length, with a denser spacing of cells along vessel walls, and more circumferential processes that cover the vessel more completely. To test whether HC mural cells are morphologically different to those in V1, we used confocal imaging of DsRed-labelled cells in fixed brain slices of mice whose vascular lumen had been filled with a fluorescent gelatin (Figure 2d). For each brain region, we calculated the distance between neighbouring mural cells’ somas, and the diameter of the vessels they occupied. Despite there being no differences in the sizes of the vessels sampled (Figure 6a), the average distance between neighbouring cells was significantly larger in HC (Figure 6b).

**Figure 6:**
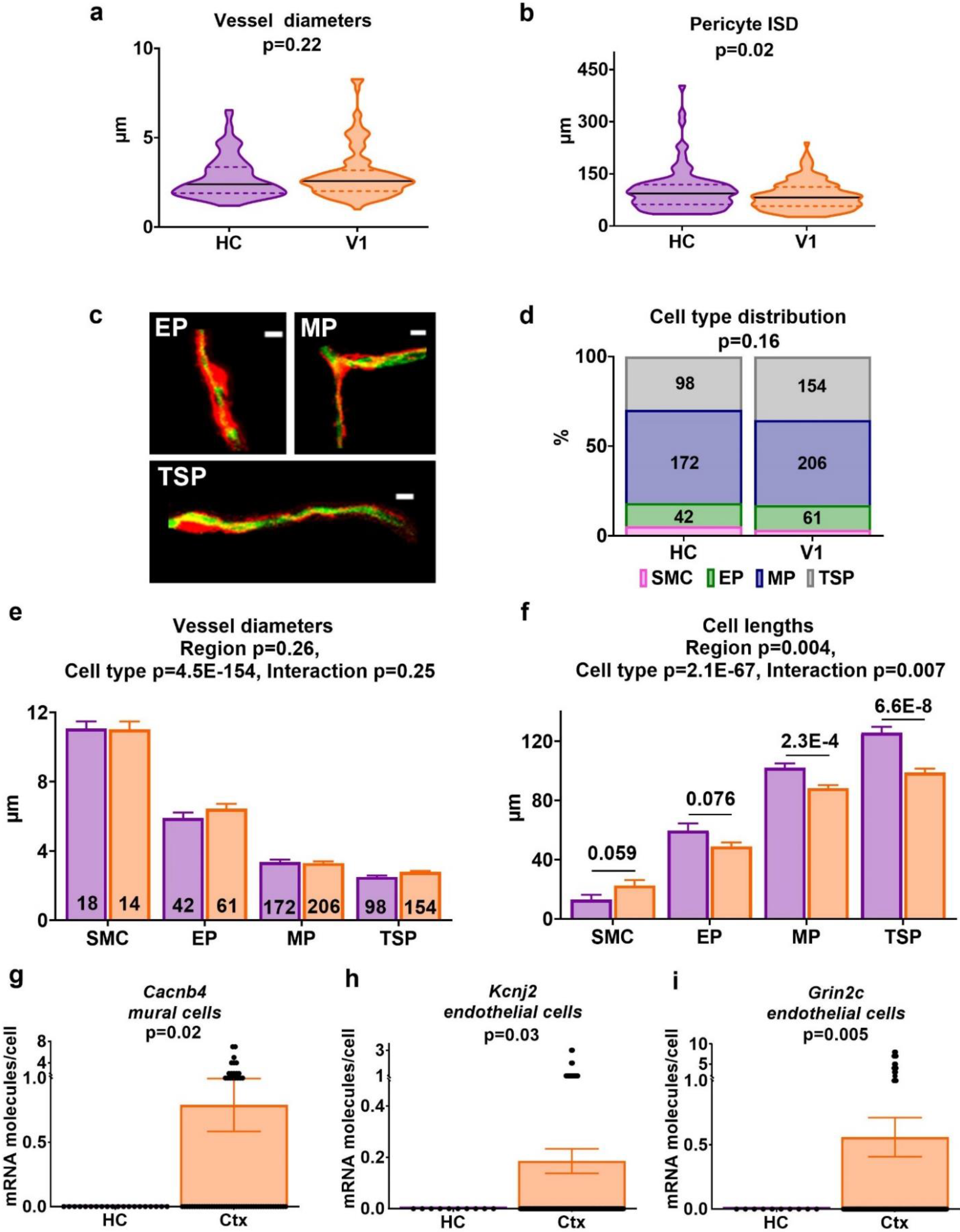
Pericyte morphology across brain regions. **(a)** Distribution of vessel diameters and **(b)** intersoma distance (ISD) between neighbouring pericytes for HC and V1 (data represent 89 vessels in HC and 127 in V1, taken from 6 mice). **(c)** Examples of ensheathing (EP), mesh (MP), and thin-strand (TSP) pericytes. Scale bars represent 5 μm. **(d)** The distribution of cell types was not different across regions (Cochran-Mantel-Haenszel 3D variant of Chi-square test). Vessel diameter **(e)** and cell length **(f)** varied with mural cell type as previously described ^12^. Diameter was not different between regions, while all the pericyte categories (EP, MP, TSP) showed a greater cell length in HC compared to V1. The numbers inside the bars show the number of vessels per each pericyte category and brain region, taken from 6 mice. There were functional differences in vascular cells between regions, as mRNA expression profiles of **(g)** Cacnb4 in cortical mural cells, and **(h)** Kcnj2 and **(i)** Grin2c in cortical endothelial cells, were higher than in HC. P-values represent the result of Welch’s t-tests, with Holm-Bonferroni corrections to correct for multiple comparisons. Dots represent individual cells (63 mural cells in cortex, 20 in HC; 126 endothelial cells in cortex, 10 in HC). P-values in graph titles are from multifactorial ANOVAs to test for effects of cell type and brain region. P-values above the bars are from post hoc t-tests with Holm-Bonferroni correction for multiple comparisons. Data in bar charts are mean +/− SEM. Horizontal lines on violin plots show median and interquartile range.

To better understand these morphological differences, we then categorised mural cells as being smooth muscle cells, or ensheathing (EP), mesh (MP) or thin-strand pericytes (TSP), based on the morphology of their processes (Figure 6c) ^12^. Smooth muscle cells form a band that encircles larger vessels. Pericytes have distinct soma and processes, with a morphology that changes along the vascular tree ^13^. Ensheathing pericytes express αSMA and are strongly contractile with a shorter length and higher vessel coverage than the less contractile mesh and thin-strand pericytes, which are sited on smaller vessels ^12^. There was no difference in the relative numbers of these different cell types between regions (Figure 6d). We next determined the length of mural cells in each category and the average diameter of the vessels which they surrounded. Our cell categorisations matched published descriptions, as vessel diameter decreased and cell length increased from ensheathing to mesh to thin-strand pericytes (Figures 6e, f). However, unlike smooth muscle cells, pericytes were longer in HC than V1. This difference was significant for mesh and thin-strand pericytes and near-significant for ensheathing pericytes (corrected p = 0.076, uncorrected p = 0.038). Because pericyte contractility is strongest nearer the cell body ^14,15^ and, generally, shorter cells are more contractile ^12^ HC pericytes may be less contractile than their counterparts in V1, which could underlie the weaker neurovascular coupling in HC.

### Functional differences between vascular cells in HC and V1 suggested by different mRNA expression profiles

In order to test for more general functional differences between vascular cells, we examined the mRNA expression profile of mural and endothelial cells using the same single-cell mRNA dataset as above ^11^ across three broad categories associated with neurovascular function: contractile machinery, ion (potassium and calcium) channels and neurovascular signalling pathways (Supplementary Tables 4 & 5). The latter category included recently identified EET and 20-HETE receptors ^16,17^ and, given the recent finding that endothelial NMDA receptors can control vascular tone ^18^, NMDA receptor subunits.

Several genes showed differential expression between V1 and HC, all of which pointed to the vasculature in V1 being more contractile or responsive than in HC (Supplementary Figure 3; Supplementary Tables 4 & 5). Mural cells showed higher expression in V1 of the calcium channel beta subunit *Cacnb4* (Figure 6g), while levels of transcripts for several other ion channel subunits were significantly higher in V1 before correction for multiple comparisons, including stargazin (*Cacng2*, which can reduce calcium channel activation ^19^), a slowly activating potassium channel (*Kcnh3*), and contractile proteins such as the skeletal muscle actin (*Acta1*) and regulators of myosin light chain phosphatase (*Ppp1r12c*) (Supplementary Table 4, Supplementary Figure 3). In endothelial cells, several more transcripts were upregulated in V1 compared to HC (Supplementary Table 5), most notably the inwardly rectifying potassium channel Kir2.1 (*Kcnj2*, Figure 6h), which has been shown to mediate the propagation of dilation from the capillary bed to upstream arterioles ^20^, the NMDA receptor subunit *Grin2c* (Figure 6i), the NO receptor *Gucy1a2* and prostaglandin E synthase (*Ptges*; Supplementary Figure 3). These transcripts would all be expected to promote dilation of the microvasculature and their lower expression in HC than V1 may mediate the weaker neurovascular function we observed physiologically. mRNA extraction from HC was not simply lower across all genes, as there was no difference between HC and V1 in the average number of mRNA molecules detected per cell across all genes investigated (Supplementary Table 6).

Thus, pericytes have a less contractile morphology in HC than V1, while HC mural and endothelial cells are less equipped to promote vasodilation via regulation of contractility, glutamate or NO sensing, prostaglandin production, or by activation of potassium channels.

### Lower oxygenation in HC may limit function

To understand how weaker HC neurovascular functioning could affect neuronal oxygen supply, and thus ATP production, we modelled oxygen diffusion and consumption in HC and V1 (Figure 7a). We first worked out how far brain tissue was, overall, from the nearest blood vessels *in vivo* (Figure 7b-c), then calculated the steady state oxygen concentration in an average capillary in each region (see Online Methods). Capillary pO_2_ was 15 mmHg (21 μM) in V1 and 10 mmHg (14 μM) in HC. Oxygen diffusion into the tissue was then simulated (Figure 7d), assuming varying rates of neuronal oxygen consumption corresponding to values reported in rodent tissue ^21,22^.

**Figure 7:**
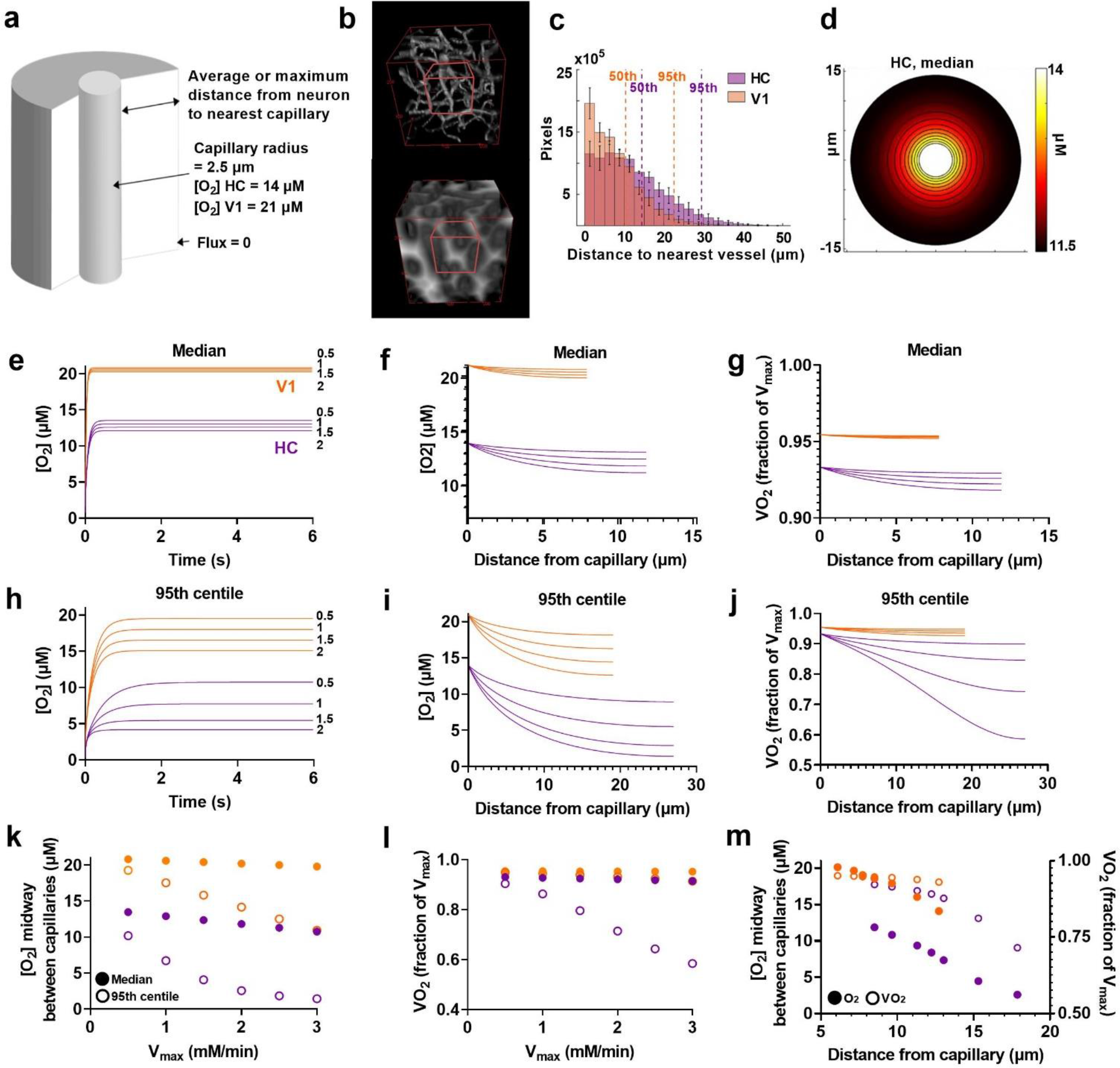
Modelling O2 diffusion in brain tissue. **(a**) Schematic of radial diffusion model. **(b)** Top: example in vivo Z-stack (250 μm^3^) of fluorescent dextran-filled vasculature in CA1 Bottom: 3D distance map generated from the z stack. White pixels denote high values and black low values. Red cube shows example 100 μm^3^ substack used to generate (c). **(c)** Average histogram of the distance of each pixel from a capillary (4 mice per region, the value for each mouse representing the average of 5 non-overlapping substacks sampled from one Z-stack). Dotted lines mark the 50^th^ and 95^th^ percentiles for HC (purple) and V1 (orange). **(d)** Example heatmap showing oxygen diffusion from a capillary in HC separated from the next capillary by twice the median tissue distance from a capillary (14.4 μm). Simulated time courses from initial conditions of zero [O_2_] for HC (purple) and V1 (orange) for tissue at **(e)** the median or **(h)** 95^th^ centile distance from a capillary. Lines represent different values of V_max_ as labelled. [O_2_] profiles across tissue for capillary separations of twice the tissue **(f)** median and **(i)** 95^th^ centile distance from a capillary. O_2_ consumption rate as a fraction of V_max_ (VO_2_) for **(g)** median and **(j)** 95^th^ centile capillary spacing conditions, calculated from oxygen profiles shown in (e) & (h). **(k)** [O_2_], and **(l)** VO_2_ reached midway between capillaries for different values of V_max_ in HC (purple) and V1 (orange) at median (solid symbols) or 95^th^ centile (hollow symbols) capillary spacings. **(m)** [O_2_] (solid symbols) and VO_2_ (hollow symbols) reached midway between capillaries for capillaries spaced at twice the 50^th^, 60^th^, 70^th^, 80^th^, 90^th^ and 95^th^ centiles (from left to right for each range of solid or hollow dots) of tissue distance from a vessel in HC (purple) and V1 (orange).

The combination of the lower capillary oxygen concentration ([O_2_]) and capillary density meant tissue oxygen levels in HC were lower than in V1 for all conditions simulated (Figures 7e,f,h,i,k). To determine if oxygen became limiting for ATP production in the tissue, we then calculated the oxygen consumption rate (VO_2_) as a proportion of the maximum rate of oxygen consumption (V_max_; Figures 7g,j,l). In V1, VO_2_ (and therefore the rate of ATP generation) occurred at over 90% of the V_max_ even far from a capillary, suggesting [O_2_] barely limited ATP synthesis. In HC, however, while VO_2_ was sustained at over 90% of the V_max_ in tissue at the median distance from a capillary, in the tissue furthest (95^th^ centile) from a capillary, [O_2_] dropped to concentrations that limited VO_2_ to under 60% of V_max_.

The effect of weak HC neurovascular coupling can be estimated by considering what happens when oxygen consumption increases but capillary oxygenation does not. Typically, neuronal activation increases net CMRO_2_ by 20-60% ^23^. In HC, increasing the V_max_ for oxygen consumption from 2 to 3 mM/min, caused [O_2_] in tissue furthest from vessels to almost halve, reducing VO_2_ from 71% to 58% of V_max_ (Figures 7k-l), while in V1, VO_2_ reduced only from 93% to 91%. Thus, the decreased SO_2_ and increased capillary spacing in HC reduce tissue [O_2_], limiting oxidative phosphorylation and ATP synthesis in the tissue furthest from a capillary, and this effect is exacerbated by weaker HC neurovascular coupling.

Finally, for a V_max_ of 2 mM/min (in the centre of the range of published values ^22^), we estimated how much of the tissue is subject to rate-limiting [O_2_] by calculating profiles for [O_2_] and VO_2_ when capillaries were separated by intermediate distances between the median and 95^th^ centile values used above (Figure 7m). This analysis suggested that low [O_2_] inhibits VO_2_ by at least 10% in 30% of HC tissue, and by at least 20% in 10% of HC tissue.

## Discussion

Our data reveal, for the first time, the properties of neurovascular coupling in mouse HC *in vivo* and demonstrate that hippocampal vascular function is compromised compared to that in neocortex in two major ways. Firstly, despite using oxygen at the same rate as the neocortex, resting blood flow in the HC is lower than in neocortex, due both to a lower vascular density and lower flow in individual capillaries. This lower energy supply combined with equivalent energy use leads the HC to have a lower resting blood oxygen saturation than neocortex. A simple model predicts that lower blood oxygenation and vascular density in the HC drive tissue oxygen to levels that readily become limiting for ATP generation. Secondly, increases in neuronal activity in the HC cause fewer and smaller dilations of local blood vessels, suggesting energy supply in the HC is less well-matched to fluctuations in energy demand than in neocortex. Our data suggest that neurovascular coupling is weaker in the HC because of differences in its vasculature rather than neuronal signalling properties, its pericytes having a less contractile morphology and its vasculature showing a pattern of mRNA expression that may be less able to promote dilation. Our model predicts that these differences in vascular physiology matter for sustaining neuronal function: the lower oxygenation and vascular density in HC interact with the decreased ability to match increased oxygen use with increases in supply. This limits tissue oxygenation in the furthest regions from blood vessels so much that ATP generation, and therefore neuronal function, are much more readily restricted in HC compared to V1.

### Lower contractility of hippocampal vasculature limits neurovascular coupling

Our conclusion that differences in microvascular function between HC and V1 underlie the weaker neurovascular coupling observed in HC stems from the observation that neuronal firing and neuronal and astrocytic expression of vasoactive messengers were no different between the brain areas, whereas differences were observed in the microvasculature. Firstly, pericytes were longer in HC than V1, suggesting functional differences that are likely to include a lower contractility, because shorter mural cells are more contractile and pericyte contractility is greatest near the soma ^12,15^. Ensheathing pericytes initiate the dilatory response to sensory stimulation in the olfactory bulb ^24^, and are amongst the earliest to respond in the neocortex ^25^. They tended to be longer in HC, so may be less contractile, helping to shape the weaker neurovascular coupling responses there. The increased length of mesh and thin strand pericytes in HC and V1 suggests they too function differently in the two regions. As observed here and previously ^14,25,26^, the smaller capillaries that contain mesh and thin strand pericytes can also dilate. Whether these dilations are active or passive remains controversial ^13,27^. However, it is likely that the longer and less contractile mesh and thin-strand pericytes in HC also contribute to weaker neurovascular coupling in HC compared to V1. Firstly, at least some of these pericytes express the contractile protein αSMA (though in a form that is less stable than in smooth muscle cells and ensheathing pericytes ^28^). Secondly, their level of intracellular calcium decreases in response to neuronal activity (consistent with a relaxation of contractile machinery ^24^). Finally, their dilations seem to mediate a large proportion of the overall increase in blood flow ^24,25^.

We also examined a published single cell RNA-Seq dataset ^11^ for differences between vascular, neuronal and astrocytic expression profiles of mRNA. The dataset comprises many more neurons and astrocytes than vascular cells, and more cortical than hippocampal cells. Thus, our power to detect differences at p = 0.05 in neuronal or astrocytic expression patterns was high (>88% with an effect size of 0.4), but much lower for endothelial (33%) or mural cells (47%). Our findings should therefore be treated with some caution and should be replicated in a dedicated experiment with larger sample sizes to verify all identified targets. Nevertheless, because *all* the positive results indicated a lack of expression in HC of proteins expected to promote dilation, despite similar levels of mRNA transcripts being detected overall in the two regions, we conclude that there is a difference in physiology between the two vascular beds.

### Weak hippocampal neurovascular coupling is not caused by the cranial window

Our use of a chronic cranial window over HC allows us to measure CBF, blood oxygenation and individual vascular and neuronal signals in a region that is normally inaccessible to two-photon imaging. It requires aspiration of a column of neocortex and insertion of a cannula through which dorsal CA1 can be imaged. We have tested whether surgery affected HC function or neurovascular coupling properties (Supplementary Figure 4). There was higher expression of inflammatory markers (GFAP and Iba1) near to the cranial window in HC than in V1, but this difference disappeared 100 μm from the window. There was no difference in neurovascular coupling properties in vessels above and below 100 μm, and spatial memory was unimpaired in HC mice. We therefore find no evidence that the more invasive HC cranial windows are driving the regional differences we observe.

### Low tissue oxygenation and weak neurovascular coupling: a perfect storm underlying HC vulnerability to disease?

Because the vascular density in HC is lower, each capillary supplies oxygen to a larger volume of tissue than in V1, so the lower blood flow we observe in individual hippocampal capillaries may be a necessary compensatory mechanism to allow increased oxygen extraction to this larger tissue volume. To our knowledge this is the first time hippocampal and neocortical oxygenation have been directly compared, and the first time hippocampal oxygenation has been measured in a sealed skull (so the tissue is not exposed to atmospheric oxygen) in awake animals.

Our estimates of capillary [O_2_] are at the lower end of previously observed values from anaesthetised preparations (14 μM in HC and 21 μM in V1), compared to 16 μM in HC ^29^ and 56 μM in cortex ^30^, which is as expected because oxygen consumption rates and pO_2_ values are lower in awake animals ^31,32^, and are within the range reported in sensory cortex of awake mice by directly measuring [O_2_] with a phosphorescent probe ^32^.

Our model predicts very different tissue [O_2_] in V1 and HC. In V1, [O_2_] gradients are very shallow, and the oxygen consumption rate by oxidative phosphorylation (i.e. the rate of ATP synthesis) is maintained at >90% of the maximum throughout the tissue. However, in HC the lower vascular density and lower oxygenation mean that [O_2_] readily becomes limiting for ATP synthesis for physiological V_max_ values (1-3 mM/min ^22^). An inhibition in the rate of oxidative phosphorylation of at least 20% likely occurs in around a tenth of HC volume. Furthermore, when oxygen use outstrips supply, oxygen consumption is barely affected in V1 but is reduced to as little as 60% of the V^max^ in 5% of HC tissue. Physiologically, HC seems adapted to deal with these conditions: it can sustain neuronal function despite oxygen levels that could be considered hypoxic. Decreases in synaptic function occur *in vitro* after induction of “hypoxic” conditions using higher [O_2_] than we estimate to be normoxic for HC (e.g. 20 mmHg/28 μM ^33^, compared to <14 μM here) but, in fact, such responses are likely to be caused by the decrease, rather than the absolute concentration of O_2_ ^34^. But while HC may cope with low [O_2_] physiologically, its lower oxygenation and weaker ability to match oxygen demand to supply can help explain its vulnerability to hypoxia and conditions where cerebral blood flow decreases, such as ischaemia and Alzheimer’s disease. Further decreases in oxygen availability caused by these conditions will produce a larger reduction in ATP synthesis over a larger volume of tissue in HC than in neocortex, because it already exists in a state where O_2_ levels are limiting. However, interventions that boost HC oxygenation may therefore prove particularly beneficial. Indeed, a drug that boosts hippocampal blood flow (nilvedipine ^35^) benefitted a subgroup of patients with mild Alzheimer’s disease ^36^, suggesting that insufficient oxygen delivery to HC might be a key early factor in Alzheimer’s disease in some individuals, and that improving oxygen delivery may be therapeutic.

### Interpreting BOLD

Our results also illuminate why BOLD signals in HC were found to be unreliably coupled to neuronal activity ^5,6^. In HC, smaller and less frequent vascular dilations in response to local changes in neuronal activity will produce smaller positive BOLD signals. BOLD signals are therefore a less sensitive measure of neuronal activity in HC than in neocortex. Simple experimental designs that compare the degree of activation across the brain could therefore erroneously conclude lower HC activation than cortex even when activity levels are the same. Analyses that test for specific patterns of activity, such as correlations of voxels with a behavioural measure, or cognitive model of interest will be less affected by the different neurovascular coupling properties in HC and V1 ^37^, but the relative insensitivity may nevertheless lead to more failures to detect subtle effects in HC than in neocortex ^38^.

Our work suggests HC physiological and pathophysiological functioning is shaped (and limited) by its vasculature, an insight that will aid understanding of disease states and human imaging studies. Further work should directly test in which situations hippocampal oxygen availability limits its function and whether boosting oxygen availability by increasing blood flow in these conditions can preserve hippocampal function. This will help guide therapeutic strategies in conditions such as Alzheimer’s disease, where boosting hippocampal blood flow in some, but not all people, can be therapeutically effective.

## Online Methods

### Animal Procedures

All experimental procedures were approved by the UK Home Office, in accordance with the 1986 Animal (Scientific Procedures) Act. All experiments used mice with a C57BL/6J background of either sex, which were either wild types (6 in total, 4 males, 2 females) or expressed GCaMP6f under the control of the Thy1 promoter (C57BL/6J-Tg(Thy1-GCaMP6f)GP5.5Dkim/J ^8^; 26 in total, 11 males, 15 females) and/or DsRed under the control of the NG2 promoter (NG2DsRedBAC ^39^; 12 total, 7 males, 5 females). Mice were housed in a 12h reverse dark/light cycle environment at a temperature of 22°C, with food and water freely available.

#### Surgery for cranial window placement

All mice used for *in vivo* imaging experiments underwent the following surgical procedure under isoflurane anaesthesia (maintained between 0.8-2%). The mouse was secured on a stereotaxic frame with a head mount (Kopf). At the beginning of the surgery mice were subcutaneously injected with 2.4 μl/g of dexamethasone (2 mg/ml), 400 μl of saline, and 1.6 μl/g of buprenorphine (0.3 mg/ml, diluted 1:10 with saline) to reduce inflammation, dehydration and post-operative pain, respectively. Temperature was maintained at 37 °C throughout using a homeothermic blanket (PhysioSuite, Kent Scientific Corporation). First, the scalp and underlying thin periosteum layer were removed across the entire dorsal skull surface, and a 3 mm circle overlaying the visual cortex or hippocampus was marked. Scratches were made in the exposed skull using a scalpel to increase the surface area for bonding with the head plate. The skin around the exposed skull was sealed using dripping tissue adhesive (3M VetBond). The mouse was then tilted on the head mount so that the area marked for the craniotomy laid flat. Black dental cement (Unifast Powder mixed with black ink (1:15 *w*/*w*) and Unifast Liquid) was applied to all areas of the exposed skull, except the marked craniotomy region and its immediate surround. A custom-made stainless steel head plate was placed over the dental cement and left for a few minutes until dry. Next, a craniotomy was performed over the previously-marked area using a dental drill (Fine Science Tools, burr size 007), after which the underlying dura was carefully removed. For visual cortex surgeries, an optical window (made from two 3 mm glass coverslips and a 5 mm glass coverslip (Harvard Apparatus) bonded together with optical adhesive (Norland Products, Inc)) was placed into the craniotomy and secured using dripping tissue adhesive and dental cement ^40^. For hippocampal surgeries, ~1.3 mm of cortex was aspirated (New Askir 30, CA-MI Srl) until the striations of the corpus callosum (just above CA1 hippocampus) were visible ^7^. A 3 mm round stainless steel cannula (2.4 mm ID, 3 mm OD, 1.5 mm height, Coopers Needle Works Ltd) with a 3 mm glass coverslip (Harvard Apparatus) attached using optical adhesive was inserted into the craniotomy and secured with dripping tissue adhesive and dental cement. The mouse was then injected with 5 μl/g of meloxicam (5 mg/ml, solution for injection, diluted 1:10 with saline) subcutaneously to further aid with post-operative pain relief. Finally, for both surgery types, 2 rubber rings were secured on top of the head plate with dental cement to allow for two-photon imaging using a water-based objective. The mouse was removed from the head mount attached to the isoflurane machine and placed into a heat box (37 °C) to recover, before being singly-housed in a recovery cage. Post-surgery meloxicam (200 μl, 1.5mg/ml) was administered orally in the food for 3 days for additional pain relief during recovery.

#### Habituation

Prior to imaging, and following a post-surgery recovery period of at least one week, mice were habituated to head-fixation on a polystyrene cylinder daily (over the course of 5 days). For the first habituation session, mice were handled by the experimenter and allowed to explore the cylinder freely without head fixation. The following sessions consisted of head-fixing the mouse atop of the cylinder for gradually increasing time periods (increasing from 1 minute in session 2 to 15 minutes in session 5).

### In vivo imaging

#### Experimental set-up

Mice were head-fixed atop of a polystyrene cylinder, fitted with a Kuebler rotary encoder (4096 steps/revolution) to measure locomotion, underneath a two-photon microscope (Scientifica) or combined laser doppler flowmetry/haemoglobin spectroscopy probe (VMS-Oxy, Moor Instruments; Figure 1a). In front of the mice were two computer screens which, when required, displayed a virtual reality maze (custom-designed in ViRMEn ^41^, MATLAB; for HC surgery mice; Figure 1c) or drifting gratings (PsychoPy, 315 ° orientation refreshed at 60 frames per second; for V1 surgery mice; Figure 1d).

#### Combined laser doppler flowmetry/haemoglobin spectroscopy (Oxy-CBF probe)

A combined laser doppler flowmetry/haemoglobin spectroscopy probe allowed for monitoring of the following net haemodynamic measures: blood flow (flux), speed, oxygen saturation (SO_2_), oxygenated haemoglobin (HbO), deoxygenated haemoglobin (Hbr), and total haemoglobin (Hbt). Experiments typically lasted 0.5-1 hours. The cerebral metabolic rate of oxygen consumption (CMRO_2_) was calculated from these variables using Equation (1) ^42^:

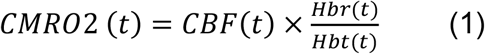

To test the reliability of the oxy-CBF probe on our rig as a chronic measure of haemodynamic activity, we ascertained that signals were stable across imaging sessions (Supplementary Figure 5a) and checked that fluorescence signals from the brain did not interfere with the haemodynamic measurements (Supplementary Figure 5) and reached zero in the absence of blood flow following the death of the subject (Supplementary Figure 5).

#### Analysis of oxy-CBF probe data

The oxy-CBF probe data (flux, speed, SO_2_, HbO, Hbr, Hbt, or CMRO_2_) was categorised as occurring during periods of rest (i.e. no locomotion and no visual screen presentations) or not. The average value of each haemodynamic parameter was calculated per animal (across sessions) during these stationary periods (Figures 2a-c).

Peaks in the CMRO_2_ trace were detected (as specified in section below: ‘*Analysis of two-photon microscopy data’*, Figures 5 e-h), and those peaks which were more than 2 standard deviations larger than their preceding baseline were extracted, along with their corresponding Hbt traces (Figures 5 f-j).

#### Two-photon microscopy

Mice were injected with 2.5% (w/v) Texas Red Dextran dissolved in saline (70 kDa via tail vein or 3 kDa subcutaneously, Sigma-Aldrich) in order to visualise blood vessels during imaging. All blood vessels recorded local to neuronal calcium activity were capillaries or arteriole branches to ensure that similar types of vessels were selected between the regions. High-resolution imaging of vessels and calcium from excitatory neurons was performed with a commercial two-photon microscope (Scientifica), a high numerical aperture water-dipping objective (20× aperture, XLUMPlanFL N, Olympus or 16× aperture, LWD, Nikon), and a Chameleon Vision II Ti:Sapphire laser (Coherent). Tissue was excited at 940nm, and the emitted light was filtered to collect red and green light from Texas Red Dextran (vessel lumen) and Thy1-GCaMP6f (excitatory neurons) respectively. Imaging sessions were recorded in SciScan software (Scientifica). Typically, imaging occurred up to a depth of ~500 μm from the surface. Imaging sessions included both wide field-of-view recordings of network neuronal calcium activity (256×256 pixels, speed range 6.10-15.26 Hz, speed average 7.75 Hz, pixel size range 1.35-2.56 μm, pixel size average 1.80 μm), and smaller field-of-view recordings of individual blood vessels and local neuronal calcium signalling (256×256 pixels, speed range 3.05-7.63 Hz, speed average 6.64 Hz; pixel size range 0.15-0.63 μm, pixel size average 0.23 μm). High speed line scans (speed range 413-2959 Hz, speed average 1092 Hz; pixel size range 0.15-0.46 μm, pixel size average 0.20 μm) were also taken from individual blood vessels to track red blood cell (RBC) velocity. Vessels and local calcium were imaged from each of the distinct layers of CA1 (stratum oriens, stratum pyramidale, stratum radiatum, stratum lacunosum-moleculare) and V1 (L1, L2/3, L4). The layers in CA1 were clearly distinguishable by the changing morphology of the pyramidal neurons over increasing depths within this region (^43^, Supplementary Figure 4a). The layers of V1 were distinguished based on the distance of the imaging site from the pial vessels ^44^. Two-photon imaging experiments typically lasted 1-3 hours.

#### Analysis of two-photon microscopy data

Excitatory calcium activity was first registered in ImageJ to correct motion artefacts, before being extracted using the CellSort package in MATLAB ^45^. CellSort identified regions of interest (ROIs) over multiple active individual cells (both soma and neuropil). Any ROIs which overlapped the same cell were either merged or removed, as appropriate. For most analyses, net activity in a recording was calculated across ROIs, except when correlations between individual ROIs were calculated (Figure 5c).

Vessel diameter (XY movies) and RBC velocity (line scans) data were registered in ImageJ to remove motion artefacts, and despeckled. The vascular lumen was bright due to the dye injection, whereas the red blood cells within the vessel and background around the vessel were dark. Custom MATLAB scripts were used to extract the diameter along the vessel branch(es), or the RBC velocity. Briefly, for each frame of XY diameter movies the diameter was calculated from the full width half maximum of the intensity perpendicular to the vessel axis at every second point along its skeleton, and averaged across a running window of 5 skeleton pixels at a time. For RBC velocity data from line scans, we adapted freely-available code ^46^. In short, the angle of the shadow cast by the RBCs travelling through the vessel was used to calculate the speed at which they travelled over successive 40ms (time, T) blocks (with an overlap of 10ms in time (i.e. T/4) between blocks).

To detect peaks in the neuronal calcium signal local to a blood vessel, the calcium trace was first normalised across the entire recording (e.g. Figure 3b) using equation (2), before identifying putative events as peaks which were >10% of the maximum. The data around each calcium peak was then extracted from the raw (unnormalized) trace (5 s before & 10 s after), and these peaks were then normalised to their own baseline (5 s preceding peak) using equation (3). Changes in calcium fluorescence from baseline (e.g. Figures 3d,j) are presented as ΔF/F. Delta is conventionally used to denote a change from the initial state, and so delta F over F compares the change of intensity after activation to that at resting baseline.

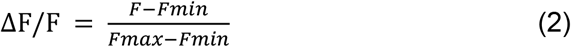

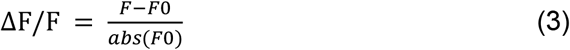

F represents the entire fluorescence trace, and F_0_ represents the baseline fluorescence period. Events were excluded if their peaks were smaller than 2 standard deviations above the mean value of the baseline. Vessel traces were extracted across the times of these calcium events, or after bootstrapping by shuffling each of the vessel traces over 100 iterations across time using the MATLAB function ‘*randperm’*, so they were no longer aligned to calcium events. For every calcium event, both the corresponding real vessel diameter trace and shuffled vessel diameter trace were classified as being “responsive” or not. Responsive diameter traces were those where dilations occurred for >0.5 s, within 5 s of the calcium event, that were greater than 1 standard deviation above baseline.

#### Behavioural testing

Animals which had previously undergone HC or V1 surgery for *in vivo* imaging underwent a simple hippocampal-dependent object location memory task to test spatial memory ^47^ (Supplementary Figure 4). The mouse was presented with two objects in a training environment for 10 minutes, before being removed from the environment. After 5 minutes in the home cage, the mouse was then returned to the environment, after one of the objects had been moved to a novel location. The time spent exploring both the familiar and novel object locations was assessed by a blinded observer hand-scoring the time per object. Any mice which spent less than 5 total seconds exploring the test environment were excluded from the analysis.

### In vitro imaging

#### Transcardial perfusion and gel-filling the vasculature

Adult mice were terminally anaesthetised with pentobarbital. A transcardial perfusion was performed using 4% paraformaldehyde (PFA) in 0.1 M phosphate buffer solution (PBS). In cases were the vasculature was labelled, this was also followed by the perfusion of 5% gelatin containing 0.2% FITC-conjugated albumin. Mice were then chilled on ice for at least 30 minutes, before brains were extracted and fixed in 4% PFA overnight. The brains were subsequently transferred to a 30% sucrose solution in PBS with 0.1% sodium azide for a minimum of 3 days, after which they were sliced (100 μm slices) on a vibratome (Leica), and stored in the fridge (at 4 °C) in PBS with 0.1% sodium azide, before immunohistochemical labelling and/or mounting.

#### Immunohistochemical labelling and slice mounting

Adult NG2-DsRed mice were used for pericyte morphology and vascular network analyses (Figures 2 & 6), as pericytes and smooth muscle cells were transgenically labelled in red. Vessel type could therefore be identified based on the smooth muscle or pericyte cell morphology.

For analysis of inflammation (Supplementary Figure 4) mice which had previously undergone HC or V1 surgery for *in vivo* imaging were used, and active astrocytes were labelled for GFAP and microglia labelled for Iba1 using immunohistochemistry. First, brain slices were washed in 1 × PBS whilst being shaken for three cycles (10 minutes per wash). The slices were then blocked in 5% normal goat serum and 0.3% Triton X-100 in 1 × PBS for an hour. Slices were incubated in the relevant primary antibodies (chicken anti-GFAP primary antibody; Abcam, ab4674, 1:500 dilution or rabbit anti-Iba1, WAKO, 019-19741, 1:600 dilution) for 36 hours at 4 °C. Slices were washed three times in PBS, before being incubated in the relevant secondary antibody for 24 hours at 4 °C (Alexa 647 goat anti-chicken, Abcam, 1:500 dilution or Alexa 647 goat anti-rabbit, Abcam, 1:500 dilution). Gel-filled and/or immunohistochemically labelled slices were washed for a final three cycles in PBS, before being mounted onto slides in Vectashield Hardset mounting medium and stored at 4 °C before imaging.

#### Confocal imaging

Imaging was performed on a confocal microscope (Leica SP8) using a 20x objective (HC PL APO CS2, 20×/0.75, dry). Continuous wave lasers with ~488, ~543, and ~633 nm excitation wavelengths were used for FITC, DsRed, and Alexa647, respectively. Images were collected with dimensions of 1024 by 1024 pixels (with a lateral resolution of 0.57 μm per pixel), averaged 4-6 times per line, and were viewed and analysed in ImageJ software.

#### Analysing pericyte and vessel morphology

For the analysis of individual pericyte cell morphology, composite z-stacks of brain slices with DsRed-labelled pericytes and FITC-albumin labelled blood vessels were used to identify the span along the vessel length of individual pericyte cells, and the vessel diameter measured at the location of the DsRed-positive cell body. The cell was then categorised as being an ensheathing, mesh or thin-strand pericyte, or a smooth muscle cell, based on its distinct morphology ^12^ (see Figure 6c for example images).

To calculate capillary density (Figures 2d, e), 2D maximum projections of FITC-labelled blood vessels were skeletonised, and the total length of the vessel skeleton within the entire region was divided by the tissue volume imaged.

#### Analysing post-surgery inflammation levels

For each GFAP- or Iba1-labelled brain slice, the z-stack was 2D maximum-projected for both the surgical and non-surgical hemisphere. In each hemisphere, three linear 500 μm ROIs were placed perpendicularly to the cranial window (or equivalent locations in the non-surgical hemisphere). The intensity profiles along these three lines were then averaged and normalised to the average value in the non-surgical hemisphere of the same slice.

### Single-cell RNA-Seq data

The raw single cell RNA-Seq data was taken from a freely-available online database ^11^ (http://linnarssonlab.org/cortex/). The authors used quantitative single-cell RNA-Seq to perform a molecular census of cells in somatosensory cortex (S1) and hippocampus (CA1) and categorise cells based on their transcriptome. We compared expression of components of neurovascular signalling pathways, ion channels, or contractile machinery in vascular cells (pericytes, smooth muscle cells, endothelial cells; Figures 6g-i, Supplementary Figure 3), and vasodilatory signalling pathways in pyramidal cells, interneurons and astrocytes (Figure 4). The levels of target transcripts in hippocampus and cortex were compared using multiple t-tests in RStudio, with a Holm-Bonferroni correction for multiple comparisons (see Supplementary Tables 1-4).

### Modelling oxygen concentrations in tissue

#### Estimating capillary oxygen concentrations

Our *in vivo* oxy-CBF probe measurements give us the saturation of oxygen (SO_2_) in the blood. From this, we calculated the partial pressure (pO_2_) in RBCs using the haemoglobin oxygen dissociation curve for C57/BL6 mice (the background of our experimental mice), with a Hill coefficient of 2.59 and a P50 of 40.2 mmHg ^48^. This yielded RBC pO_2_ values of 30 mmHg (42 μM at 37 °C) in HC and 43 mmHg (60 μM) in V1. (Micromolar concentrations of oxygen in tissue at 37 °C were calculated from partial pressures using the Henry’s Law-based conversions freely-provided by PreSens Precision Sensing; https://www.presens.de/support-services/download-center/tools-utilities.html). Our calculated RBC oxygen levels were consistent with those measured with a phosphorescent probe in somatosensory cortical and olfactory bulb RBCs in an awake mouse (30-100 mmHg) ^32,49^, in experiments that also measured pO_2_ between RBCs (interRBC pO_2_). We then estimated the interRBC pO_2_ for our experiments by using those interRBC phosphorescent probe measurements which had the same pO_2_ as we observed ^32,49^. This yielded an estimate of interRBC [O_2_] in V1 of 15 mmHg/ 21 μM and in HC of 10 mmHg/ 14 μM. Because tissue pO_2_ equilibrates with interRBC pO_2_ rather than RBC pO_2_ ^49^, we used these estimates of interRBC [O_2_] as the capillary [O_2_] in our model.

#### Calculating capillary spacings

Using ImageJ (Exact Euclidean Distance Transform (3D) plugin) we calculated a 3D distance map from a smoothed and binarized *in vivo* z stack of the vascular network in each brain region. We extracted 5 substacks of 100×100×100 μm from each distance map. The distribution of the distances of pixels from the nearest vessel were then extracted and averaged across all substacks from a given larger z stack. Histograms of these data were averaged across 5 stacks per brain region (Figure 7c) and the 50^th^, 95^th^ and intermediate centile distances from a capillary were calculated (Figures 7e-l, m). For a given bar in the histogram, contributing pixels are either at a midpoint between two capillaries, or are closer to one capillary than another. Those pixels above the 95^th^ centile must lie more-or-less at the midpoint between vessels, while many pixels at the median distance from the vessel will be nearer to one vessel than another. However [O_2_] at each distance from a vessel would be larger if that point is at the midpoint between two vessels compared to be being nearer to one vessel than another (and therefore oxygen from that vessel is having to feed a larger tissue volume). For the diffusion model below, we therefore used the distribution of tissue distances from a capillary as a conservative estimate of capillary separation, meaning that our estimates of oxygen gradients are, if anything, slightly overestimated.

#### Diffusion model

We used the Heat Transfer functions in the Partial Differential Equation Toolbox of MATLAB to numerically solve Fick’s diffusion equation for radial geometries with Michaelis-Menten consumption of oxygen through oxidative phosphorylation (Equation (4)):

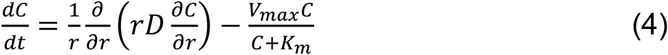

where *r* is the distance from the centre of a capillary (of radius 2.5 μm; Figure 2g), *C* is the concentration of O_2_, *D* is its diffusion coefficient in brain at 37 °C (9.24 × 10^−9^ /min ^50^), *K_m_* = 1 μM is the EC_50_ for O_2_ activating oxidative phosphorylation ^51^, and *V*_*max*_ is the maximum rate of oxidative phosphorylation at saturating [O_2_]. The initial conditions were C(r) = 0, and the boundary conditions fixed [O_2_] at the edge of the capillary (*r*(2.5)) to be the calculated interRBC [O_2_] for each brain region and at the midway point between two capillaries (r(max) *dC*/*dr* = 0). The equation was solved over values for r from 2.5 μm up to the average tissue distances from a capillary as calculated above (median, 95^th^ or intermediate centiles) for each brain region.

#### Oxygen consumption

Values for *V*_*max*_ (0.5-3 mM/min) were chosen based on the range of brain oxygen consumption rates previously reported ^22^. Of this range, values of 1-2 mM/min may be the most physiological, being reported from recent experiments measuring the oxygen gradient using phosphorescent lifetime imaging ^e.g. 21^ (rather than an invasive electrode) and matching the whole brain averaged oxygen consumption rate ^52,53^. However, previous measurements in rodents were done under anaesthesia, so rates may be higher in our awake system.

### Statistical analysis

All data are presented as mean +/− SEM or individual data points. P-values are from ANOVAs (one-way univariate or multifactorial), Bonferroni post-hoc comparison tests, independent sample Student t-tests, Welch’s independent sample t-tests, chi-square tests (2×2 contingency with Fisher’s exact significance test, or 3D Cochran–Mantel–Haenszel (CMH) tests with Pearson’s R multiple post-hoc comparisons) as stated. Welch’s t-tests are used instead of Student’s t-tests in cases of unequal variance between groups. In the cases of multiple comparisons, a procedure equivalent to the Holm-Bonferroni correction was applied to calculate an adjusted p-value. This correction consisted of ranking the p-values outputted from t-tests in ascending order, and then multiplying the lowest p value by the number of comparisons, the second lowest p value by the number of comparisons minus one, and so on, until comparisons were no longer significant at p = 0.05. Statistical analyses were conducted using SPSS, RStudio, or MATLAB, and figures were created using GraphPad Prism.

## Supporting information

Statistics Reports

## Author Contributions

K.S, L.B, K.B, D.M.G, and D.C collected data for the studies. K.S, H.C and C.N.H designed and analysed the studies. K.S, D.M.G, O.B and C.N.H wrote scripts which contributed to the analysis of the data. K.S and C.N.H wrote the manuscript with feedback from all authors.

## Acknowledgements

This work was supported by an MRC Discovery Award, an Academy of Medical Sciences/Wellcome Trust Springboard Award to C.N.H., the Alzheimer’s Society for O.B.’s studentship and Sussex Neuroscience PhD studentships for D.C. and D.M.G, Sussex University Research Development funding to H.C. and C.N.H. and School of Psychology funding to C.N.H.

