## Supplementary Figures and Tables for "Hippocampus has lower oxygenation and weaker control of brain blood flow than cortex, due to microvascular differences"

Shaw et al.

#### Supplementary Figures

##### Supplementary Figure 1

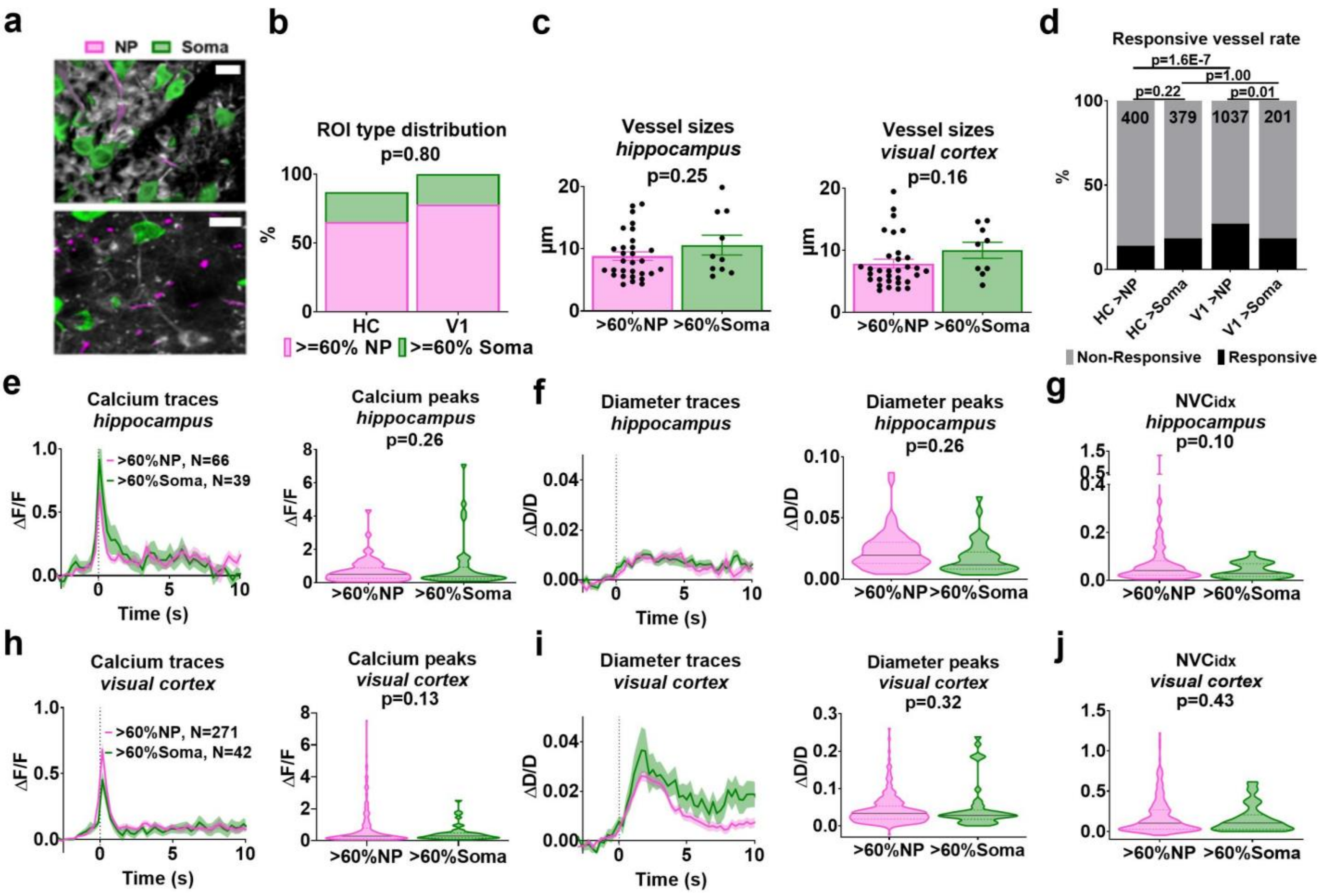


**Supplementary Figure 1: The contribution of cellular input to vascular responses**

**(a)** GCaMP6f-positive pyramidal neurons (white) with regions of interest (ROIs) categorised as soma (green) or neuropil (NP, pink) based on their morphology, for HC (top) and V1 (bottom). Scale bars represent 10 µm. **(b)** Neuronal calcium activity from individual recordings was classified by whether ROIs were majority NP or majority soma (>60% of the total ROIs; V1, N=313 calcium events, 7 mice; HC, N=105 calcium events, 6 mice; 15 recordings were excluded as there was no majority for either NP or soma). P value is from a 2x2 Chi-square contingency test. **(c)** Vessel sizes recorded in HC and V1 were the same for NP and soma recordings. **(d)** Response frequencies compared across brain regions, split by ROI type using the Cochran-Mantel-Haenszel 3D variant of a Chi-square test. Vessels near to NP were significantly more likely to dilate compared to those near somas in V1. Average traces and peak responses of **(e)** calcium, and **(f)** corresponding vessels in HC were not different if calcium signals came from NP or soma. **(g)** $NVC_{idx}$ was not different in HC depending on neuronal ROI type. Average traces and peak responses for **(h)** calcium and **(i)** corresponding vessels in V1 were not different if calcium signals came from NP or soma. **(j)** $NVC_{idx}$ was not different in V1 depending on neuronal ROI type. P-values are from independent sample t-tests, unless otherwise stated. Multifactorial ANOVAs also revealed that the regional differences in diameter responses and NVC indices were not affected by ROI type (Statistics Report Table 5d).

**Supplementary Figure 2**

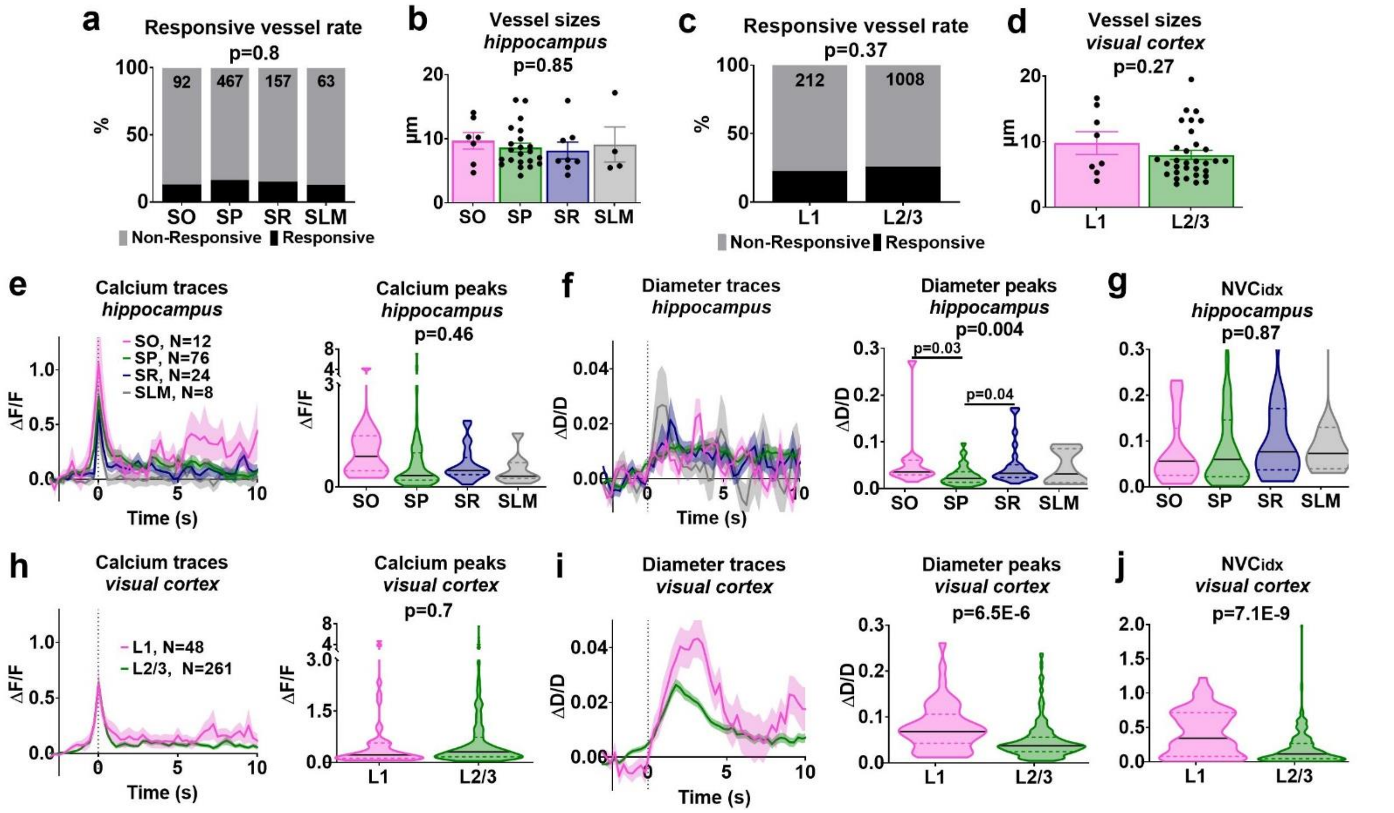


**Supplementary Figure 2: The contribution of laminar organisation to vascular responses**

Vessel recordings were separated by layer for HC (N=120 calcium events, 6 mice; SO = stratum oriens, SP = stratum pyramidale, SR = stratum radiatum, SLM = stratum lacunosum-moleculare) and V1 (N=309 calcium events, data from L4 was excluded due to low sample sizes (4 responsive events), 7 mice). Response frequencies were compared across layers for **(a)** HC (using the Cochran-Mantel-Haenszel 3D variant of a Chi-square test), and **(c)** V1 (using a 2x2 Chi-Square test). The vessel sizes sampled for each layer of **(b)** HC and **(d)** V1. The average traces and their maximum peaks for **(e)** calcium, and **(f)** corresponding vessels in HC split by layer. Vessel dilations were smallest in layer SP. **(g)** The $NVC_{idx}$ did not differ by layer in HC. The average traces and their maximum peaks for **(h)** calcium and **(i)** corresponding vessels in V1, when split by layer. Vessel dilations were largest in L1. **(j)** The $NVC_{idx}$ was largest in L1, due to bigger vessel dilations. $NVC_{idx}$ in both L1 and L2/3 were nevertheless larger than in HC (one-way ANOVA: L1, L2/3 and average across HC; Statistics report 6i). Unless otherwise stated, p-values represent the result of independent sample t-tests in V1, and one-way ANOVAs with Bonferroni post-hoc comparisons in HC.

#### Supplementary Figure 3

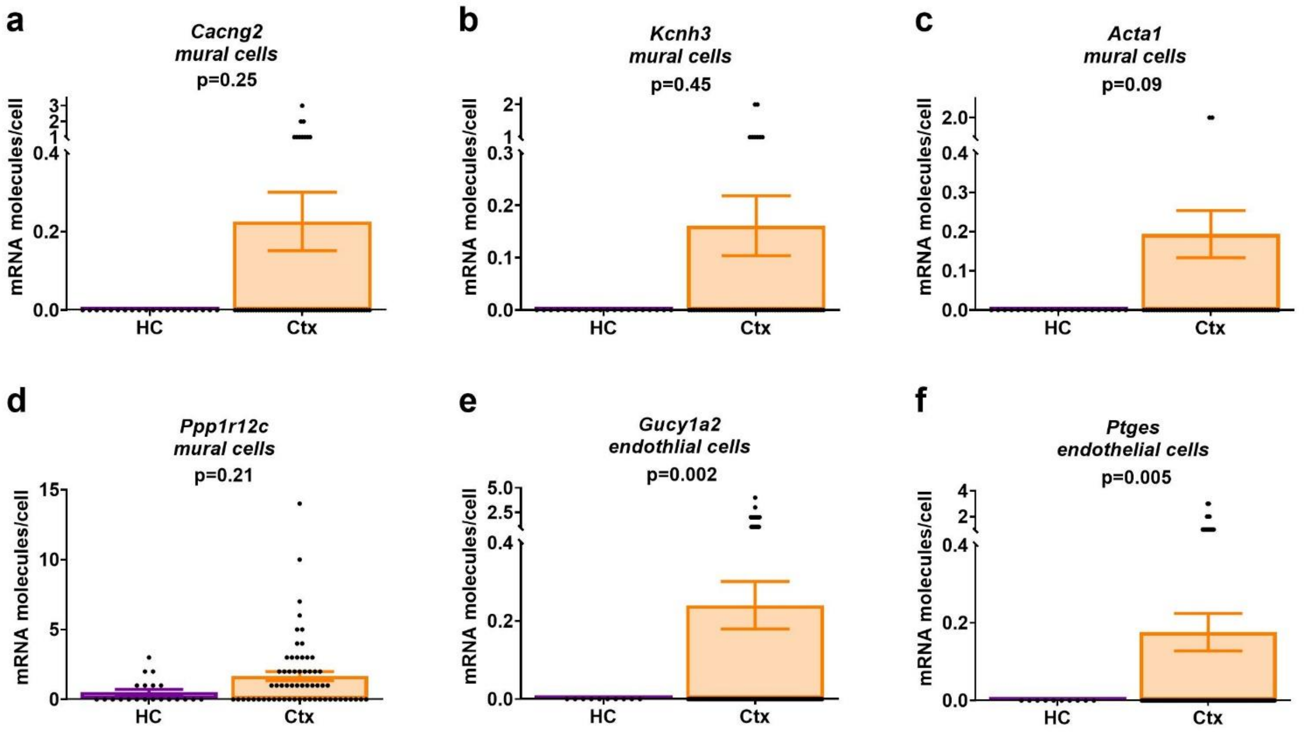


##### Supplementary Figure 3: Neurovascular pathways in vascular cells

The mRNA expression profile of mural and endothelial cells was tested across three broad categories known to link neurovascular function: ion channels, contractile machinery and neurovascular signalling pathways. Regions showed significantly different expression levels of **(a)** *Cacng2***, (b)** *Kcnh3*, **(c)** *Acta1*, **(d)** *Ppp1r12c*, **(e)** *Gucy1a2* and **(f)** *Ptges*. Dots represent individual cells. P-values represent the result of independent sample t-tests, with Holm-Bonferroni corrections for the multiple comparisons presented in each of Supplementary Data Tables 4-5.

#### Supplementary Figure 4

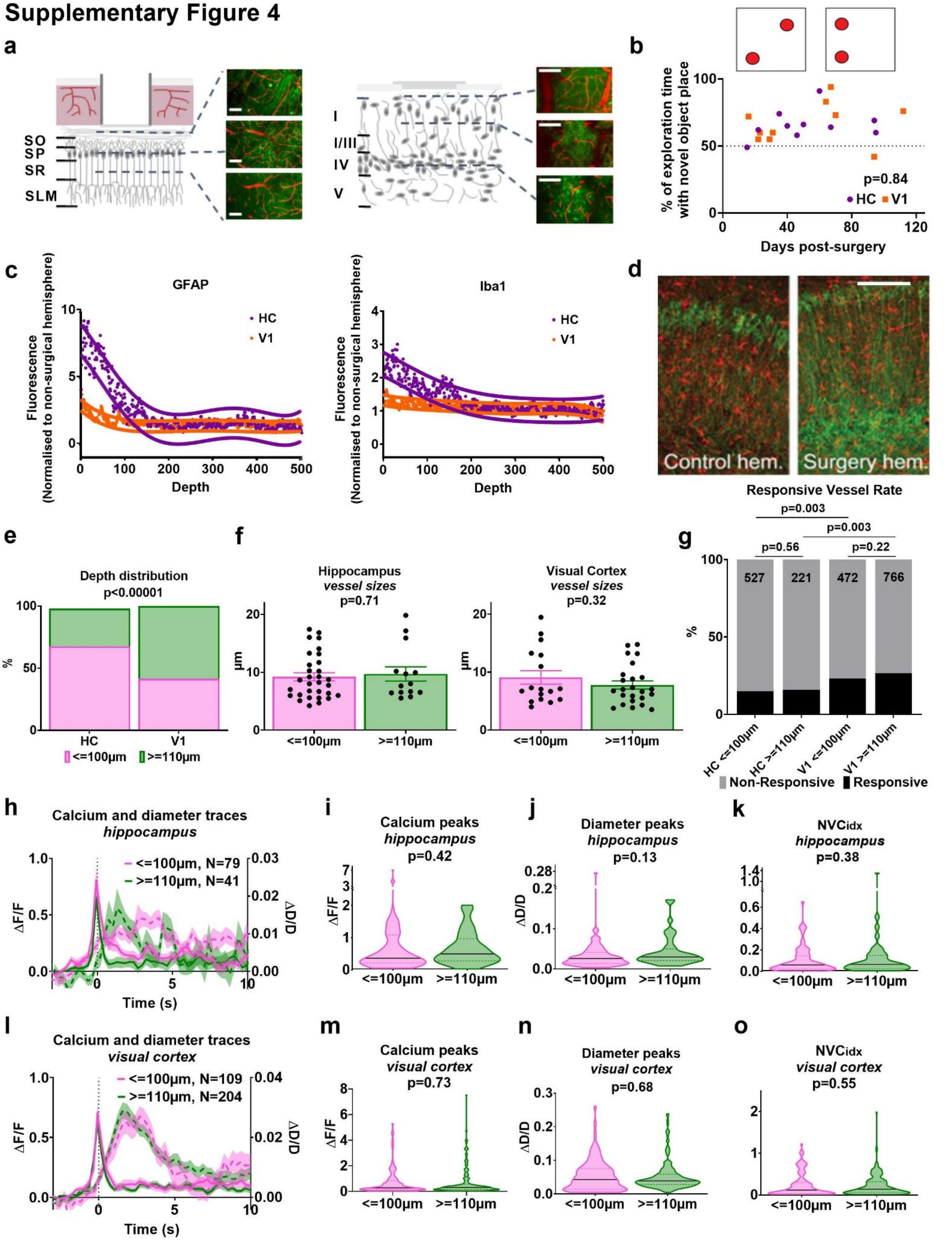


**Supplementary Figure 4: The impact of hippocampal cranial window surgery**

**(a)** Schematic showing the GCaMP6f-positive pyramidal neurons (green) and blood vessels (red) accessible for two-photon imaging after hippocampal (left) or visual cortex (right) surgery, with example maximum-projected images across layers. Scalebars represent 100µm. **(b)** Memory was assessed on a hippocampal-dependent novel object location task for mice which had undergone HC or V1 surgery. **(c)**

Average profiles for intensity of GFAP (left: HC, N=2 mice; V1, N=3 mice) and Iba1 labelling (right: HC, N=3 mice, V1, N=3 mice). Data from each mouse is an average of inflammation profiles from 3-5 slices. Exponential fits to the data with 95% upper and lower confidence intervals are plotted. Inflammation levels were considered not to be different at the point where confidence intervals overlapped (Iba1: 105 µm, GFAP: 110 µm). **(d)** Example images showing GCaMP6f-labelled neurons (green) and Iba1-labelled microglia (red) in the surgical and non-surgical hemispheres. Data above and below 100 µm from the window was compared. **(e)** Number and **(f)** sizes of vessels at different depths. **(g)** Response frequencies were not different across imaging depth but were different between brain regions (Cochran-Mantel-Haenszel 3D variant of a Chi-square test). The average traces for calcium (solid lines) and vessel diameter (dotted lines) **(h**: HC**; l**: V1**).** The corresponding maximum peak values for calcium and diameter traces **(i**: HC calcium**, j**: HC diameter; **m**: V1 calcium, **n**: V1 diameter**)**, and the $NVC_{idx}$ **(k**: HC**, o**: V1**)**, showed no differences based on imaging depth. HC: N=120 calcium events, 6 mice; V1: N=313 calcium events, 7 mice. P-values represent the result of independent sample t-tests, unless stated.

### Supplementary Figure 5

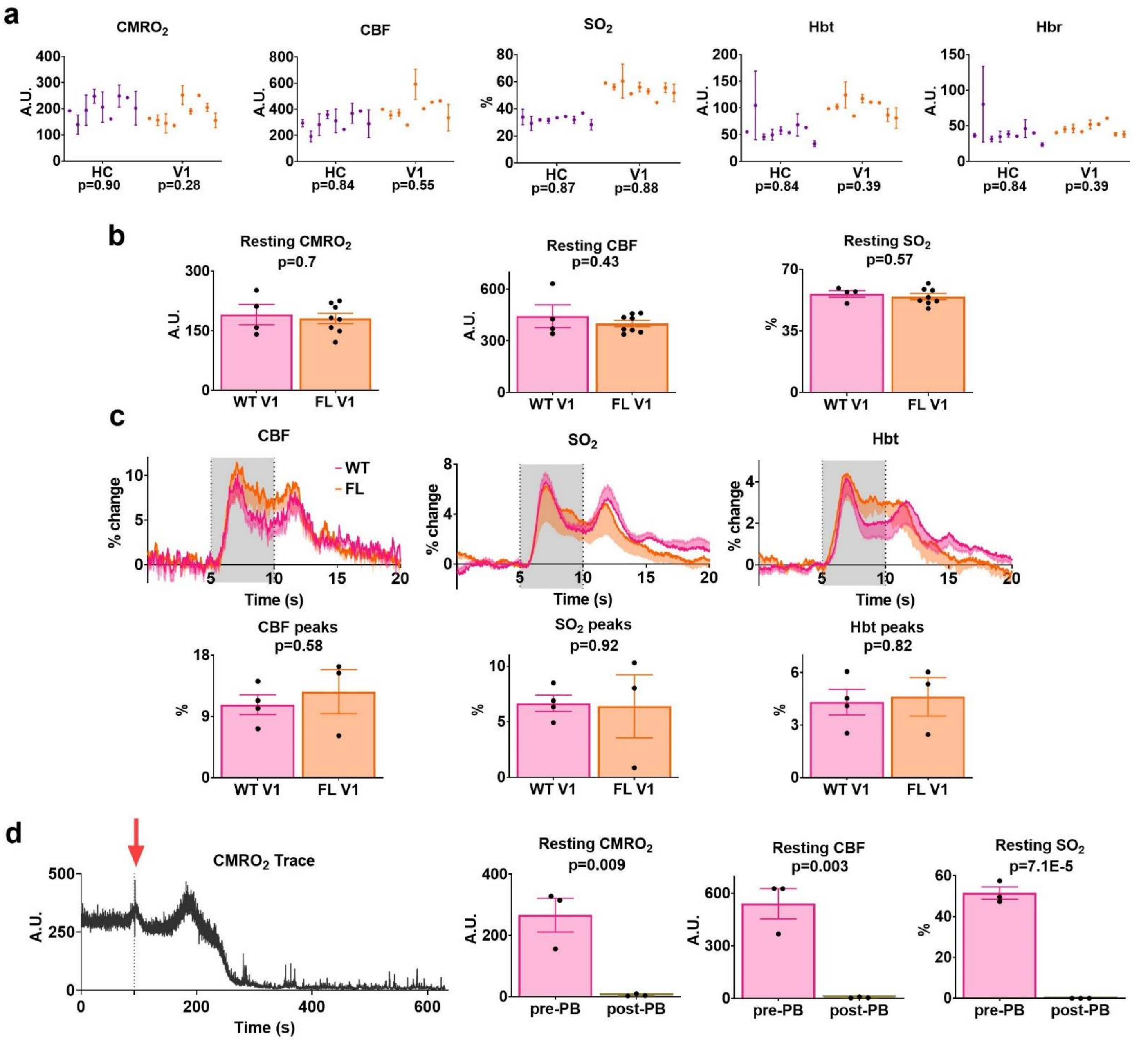


#### Supplementary Figure 5: Testing the reliability of oxy-CBF probe measurements

**(a)** The average value (mean +/- SEM) for each animal included in Figures 2a-c across the haemodynamic parameters measured by the oxy-CBF probe for $CMRO_2$, flux, $SO_2$, Hbt, and Hbr. P-values below the graphs represent the results of one-way ANOVA tests for effects of animal ID on haemodynamics. **(b)** Resting $CMRO_2$, blood flow and $SO_2$ levels were compared for GCaMP6f-positive V1 mice (N=8 mice, taken from Figures 2a-c) to wild type V1 mice (non-fluorescent, N=4 mice across 12 sessions) to check for any effects of brain fluorescence on oxy-CBF measures. No significant differences were observed. **(c)** Stimulus-induced haemodynamic responses were also compared between wild type and GCaMP6f-positive V1 mice. **(d)** Wild-type mice were injected with pentobarbital (i.p.) while recording with the oxy-CBF probe to determine the baseline for the various parameters when blood flow ceased (N=3 mice). The black trace on the left of the bar plots shows an example continuous recording of $CMRO_2$, with the red arrow and dotted line representing when the injection was administered. Resting levels of $CMRO_2$, CBF and $SO_2$ were averaged during rest periods before the injection (i.e. alive), and for the last one minute of the recording for comparison (i.e. after death).

#### Supplementary Tables

##### Supplementary Tables 1-5: Single Cell RNA-Seq analyses

Data were taken from Zeisel et al [3] to allow comparison of mRNA transcript expression in neural, astrocytic and vascular cells of cortex and CA1 of hippocampus. The number of transcripts per cell were compared using Welch's t-tests, adjusted for multiple comparisons using a method equivalent to the Holm-Bonferroni method (for N comparisons, the most significant p-value is multiplied by N, the 2nd most significant by N-1, the 3rd most significant by N-2, etc.; corrected p-values are deemed significant if they are less than 0.05). Numbers of each cell type were as follows:

Pyramidal cells. Cortex: 398; HC: 941
Interneurons. Cortex: 169; HC: 126
Astrocytes. Cortex: 143; HC: 80
Mural cells. Cortex: 63; HC: 20
Endothelial cells. Cortex: 126; HC: 10

For neurons and astrocytes, mRNA transcripts were studied that code for neurovascular signalling molecules. For mural cells (smooth muscle cells and pericytes) we probed for contractile machinery, ion channels expected to modulate dilation ($K^+$ and $Ca^{2+}$ channels) and receptors or synthetic enzymes for neurovascular signalling molecules.

| **Supplementary Tables Key: (SD = standard deviation)** |
|---|
| Significant after correction for multiple comparisons |
| Significant before correction for multiple comparisons |
| Non-significant |

##### Supplementary Table 1: Single cell mRNA comparisons for pyramidal cells

**See Figures 4a-b**

| Gene name | Cortex Mean | Cortex SD | HC Mean | HC SD | p-value | Adj. p-value | Effect Size |
|---|---|---|---|---|---|---|---|
| ***Nos1*** | 0.005 | 0.071 | 0.053 | 0.238 | 0.000 | <0.001 | 0.237 |
| ***Ptges3*** | 2.422 | 2.842 | 3.077 | 2.858 | 0.000 | 0.001 | 0.229 |
| ***Ptgs1*** | 0.118 | 0.539 | 0.045 | 0.245 | 0.010 | 0.086 | 0.205 |
| ***Pla2g4a*** | 0.058 | 0.282 | 0.021 | 0.178 | 0.017 | 0.139 | 0.170 |
| ***Ptgs2*** | 0.261 | 0.763 | 0.163 | 0.577 | 0.021 | 0.149 | 0.154 |
| ***Ptges*** | 0.048 | 0.326 | 0.019 | 0.159 | 0.096 | 0.575 | 0.129 |
| ***Ptges3l*** | 0.168 | 0.545 | 0.121 | 0.364 | 0.115 | 0.577 | 0.110 |
| ***Pld2*** | 0.050 | 0.251 | 0.071 | 0.285 | 0.178 | 0.711 | 0.077 |
| ***Ptges2*** | 0.405 | 0.677 | 0.439 | 0.732 | 0.410 | 1.000 | 0.048 |
| ***Pla2r1*** | 0.013 | 0.132 | 0.010 | 0.097 | 0.685 | 1.000 | 0.027 |
| ***Pld1*** | 0.035 | 0.198 | 0.026 | 0.171 | 0.398 | 1.000 | 0.054 |

##### Supplementary Table 2: Single cell mRNA comparisons for interneurons

| Gene name | Cortex Mean | Cortex SD | HC Mean | HC SD | p-value | Adj. p-value | Effect Size |
|---|---|---|---|---|---|---|---|
| ***Ptges*** | 0.054 | 0.274 | 0.008 | 0.089 | 0.044 | 0.488 | 0.212 |
| ***Ptges2*** | 0.411 | 0.686 | 0.595 | 0.887 | 0.054 | 0.536 | 0.237 |
| ***Nos1*** | 0.065 | 0.331 | 0.040 | 0.196 | 0.405 | 1.000 | 0.092 |
| ***Ptges3*** | 3.911 | 4.044 | 3.794 | 3.693 | 0.796 | 1.000 | 0.030 |
| ***Ptges3l*** | 0.054 | 0.367 | 0.032 | 0.176 | 0.501 | 1.000 | 0.073 |
| ***Pla2r1*** | 0.006 | 0.077 | 0.000 | 0.000 | 0.319 | 1.000 | 0.102 |
| ***Pld1*** | 0.030 | 0.203 | 0.008 | 0.089 | 0.214 | 1.000 | 0.133 |
| ***Pld2*** | 0.155 | 0.424 | 0.111 | 0.316 | 0.312 | 1.000 | 0.115 |
| ***Ptgs1*** | 0.220 | 0.687 | 0.341 | 0.647 | 0.123 | 1.000 | 0.181 |
| ***Ptgs2*** | 0.012 | 0.109 | 0.000 | 0.000 | 0.158 | 1.000 | 0.145 |
| ***Pla2g4a*** | 0.030 | 0.230 | 0.032 | 0.217 | 0.940 | 1.000 | 0.009 |

##### Supplementary Table 3: Single cell mRNA comparisons for astrocytes

**See Figure 4c**

| Gene name | Cortex Mean | Cortex SD | HC Mean | HC SD | p-value | Adj. p-value | Effect Size |
|---|---|---|---|---|---|---|---|
| ***Ptges3*** | 0.483 | 0.786 | 1.513 | 2.882 | 0.002 | 0.040 | 0.561 |
| ***Ptgs1*** | 0.259 | 0.590 | 0.100 | 0.341 | 0.012 | 0.186 | 0.308 |
| ***Cyp2j6*** | 0.930 | 1.727 | 0.525 | 1.125 | 0.036 | 0.533 | 0.263 |
| ***Pld1*** | 0.070 | 0.328 | 0.013 | 0.112 | 0.058 | 0.818 | 0.212 |
| ***Ptges*** | 0.238 | 0.650 | 0.113 | 0.450 | 0.092 | 1.000 | 0.214 |
| ***Ptges2*** | 0.119 | 0.384 | 0.213 | 0.469 | 0.130 | 1.000 | 0.225 |
| ***Ptges3l*** | 0.042 | 0.201 | 0.025 | 0.224 | 0.574 | 1.000 | 0.081 |
| ***Pla2r1*** | 0.021 | 0.186 | 0.000 | 0.000 | 0.181 | 1.000 | 0.140 |
| ***Pld2*** | 0.329 | 0.700 | 0.413 | 0.706 | 0.395 | 1.000 | 0.119 |
| ***Ptgs2*** | 0.014 | 0.118 | 0.025 | 0.157 | 0.585 | 1.000 | 0.083 |
| ***Pla2g4a*** | 0.070 | 0.422 | 0.038 | 0.249 | 0.471 | 1.000 | 0.088 |
| ***Cyp2c68*** | 0.000 | 0.000 | 0.013 | 0.112 | 0.320 | 1.000 | 0.187 |
| ***Cyp2j5*** | 0.000 | 0.000 | 0.013 | 0.112 | 0.320 | 1.000 | 0.187 |
| ***Cyp2j9*** | 1.084 | 1.714 | 1.163 | 1.642 | 0.736 | 1.000 | 0.047 |
| ***Cyp2j11*** | 0.007 | 0.084 | 0.000 | 0.000 | 0.319 | 1.000 | 0.104 |
| ***Cyp2j12*** | 0.021 | 0.186 | 0.000 | 0.000 | 0.181 | 1.000 | 0.140 |
| ***Cyp2j13*** | 0.014 | 0.118 | 0.000 | 0.000 | 0.158 | 1.000 | 0.148 |

#### Supplementary Table 4: Single cell mRNA comparisons for mural cells

a. **Contractile machinery:**

**See Supplementary Figure 3a-b**

| Gene name | Cortex Mean | Cortex SD | HC Mean | HC SD | p-value | Adj. p-value | Effect Size |
|---|---|---|---|---|---|---|---|
| ***Acta1*** | 0.194 | 0.474 | 0.000 | 0.194 | 0.002 | 0.099 | 0.468 |
| ***Ppp1r12c*** | 1.661 | 2.541 | 0.550 | 1.111 | 0.004 | 0.206 | 0.492 |
| ***Ppp1r9a*** | 0.500 | 1.004 | 0.150 | 0.350 | 0.024 | 1.000 | 0.391 |
| ***Ppp1r13b*** | 0.161 | 0.606 | 0.000 | 0.161 | 0.040 | 1.000 | 0.305 |
| ***Calm2*** | 13.194 | 17.675 | 7.450 | 5.744 | 0.047 | 1.000 | 0.361 |
| ***Ppp1r1b*** | 0.081 | 0.329 | 0.000 | 0.081 | 0.058 | 1.000 | 0.281 |
| ***Ppp1r26*** | 0.081 | 0.375 | 0.000 | 0.081 | 0.096 | 1.000 | 0.246 |
| ***Acta2*** | 43.839 | 40.397 | 30.500 | 13.339 | 0.120 | 1.000 | 0.350 |
| ***Des*** | 0.581 | 0.984 | 1.000 | -0.419 | 0.146 | 1.000 | 0.411 |
| ***Tpm3*** | 0.613 | 1.077 | 1.200 | -0.587 | 0.153 | 1.000 | 0.472 |
| ***Calm1*** | 17.597 | 28.506 | 12.050 | 5.547 | 0.162 | 1.000 | 0.221 |
| ***Ppp1r11*** | 0.532 | 1.251 | 0.250 | 0.282 | 0.164 | 1.000 | 0.251 |
| ***Ppp1r3b*** | 0.048 | 0.282 | 0.000 | 0.048 | 0.182 | 1.000 | 0.196 |
| ***Ppp1r3g*** | 0.048 | 0.282 | 0.000 | 0.048 | 0.182 | 1.000 | 0.196 |
| ***Ppp1r16b*** | 0.371 | 0.707 | 0.200 | 0.171 | 0.188 | 1.000 | 0.264 |
| ***Ppp1r2*** | 0.597 | 1.273 | 0.350 | 0.247 | 0.240 | 1.000 | 0.215 |
| ***Ppp1r18*** | 0.016 | 0.127 | 0.100 | -0.084 | 0.249 | 1.000 | 0.450 |
| ***Calm3*** | 1.323 | 2.209 | 0.900 | 0.423 | 0.274 | 1.000 | 0.210 |
| ***Ppp1r3f*** | 0.048 | 0.381 | 0.000 | 0.048 | 0.321 | 1.000 | 0.145 |
| ***Ppp1r36*** | 0.016 | 0.127 | 0.000 | 0.016 | 0.321 | 1.000 | 0.145 |
| ***Ppp1r16a*** | 0.016 | 0.127 | 0.000 | 0.016 | 0.321 | 1.000 | 0.145 |
| ***Ppp1r13l*** | 0.016 | 0.127 | 0.000 | 0.016 | 0.321 | 1.000 | 0.145 |
| ***Mylk*** | 2.677 | 3.018 | 3.400 | -0.723 | 0.343 | 1.000 | 0.242 |
| ***Ppp1ca*** | 0.952 | 1.431 | 0.700 | 0.252 | 0.410 | 1.000 | 0.186 |
| ***Ppp1r12b*** | 0.081 | 0.329 | 0.200 | -0.119 | 0.467 | 1.000 | 0.269 |
| ***Ppp1r15a*** | 2.306 | 3.687 | 2.800 | -0.494 | 0.467 | 1.000 | 0.146 |
| ***Vim*** | 4.565 | 4.762 | 5.450 | -0.885 | 0.512 | 1.000 | 0.181 |
| ***Ppp1r10*** | 0.161 | 0.413 | 0.250 | -0.089 | 0.513 | 1.000 | 0.198 |
| ***Ppp1r35*** | 0.016 | 0.127 | 0.050 | -0.034 | 0.525 | 1.000 | 0.218 |
| ***Tpm4*** | 1.484 | 1.725 | 1.700 | -0.216 | 0.539 | 1.000 | 0.133 |
| ***Ppp1r14c*** | 0.065 | 0.248 | 0.100 | -0.035 | 0.643 | 1.000 | 0.135 |
| ***Ppp1r7*** | 0.306 | 0.692 | 0.250 | 0.056 | 0.672 | 1.000 | 0.088 |
| ***Tpm1*** | 9.774 | 8.360 | 10.700 | -0.926 | 0.683 | 1.000 | 0.109 |
| ***Ppp1r14b*** | 0.113 | 0.367 | 0.150 | -0.037 | 0.696 | 1.000 | 0.101 |
| ***Ppp1cb*** | 1.500 | 1.990 | 1.650 | -0.150 | 0.734 | 1.000 | 0.079 |
| ***Ppp1r3d*** | 0.065 | 0.307 | 0.100 | -0.035 | 0.744 | 1.000 | 0.103 |
| ***Ppp1r21*** | 0.081 | 0.275 | 0.100 | -0.019 | 0.804 | 1.000 | 0.068 |
| ***Ppp1cc*** | 0.226 | 0.612 | 0.250 | -0.024 | 0.849 | 1.000 | 0.042 |
| ***Ppp1r9b*** | 0.177 | 0.426 | 0.200 | -0.023 | 0.862 | 1.000 | 0.050 |
| ***Ppp1r15b*** | 0.274 | 0.705 | 0.250 | 0.024 | 0.874 | 1.000 | 0.036 |
| ***Ppp1r3c*** | 0.323 | 1.021 | 0.350 | -0.027 | 0.891 | 1.000 | 0.029 |
| ***Ppp1r1a*** | 0.113 | 0.367 | 0.100 | 0.013 | 0.908 | 1.000 | 0.033 |
| ***Ppp1r14a*** | 0.210 | 0.517 | 0.200 | 0.010 | 0.932 | 1.000 | 0.020 |

| | | | | | | | |
|---|---|---|---|---|---|---|---|
| ***Ppp1r12a*** | 2.387 | 4.006 | 2.450 | -0.063 | 0.935 | 1.000 | 0.017 |
| ***Myh11*** | 5.806 | 6.633 | 5.700 | 0.106 | 0.941 | 1.000 | 0.017 |
| ***Tpm2*** | 7.984 | 8.818 | 7.850 | 0.134 | 0.952 | 1.000 | 0.015 |
| ***Ppp1r8*** | 0.097 | 0.534 | 0.100 | -0.003 | 0.973 | 1.000 | 0.007 |
| ***Ppp1r2.ps3*** | 0.048 | 0.216 | 0.050 | -0.002 | 0.978 | 1.000 | 0.007 |

**b. Ion channels:**

**See Figure 6g, and Supplementary Figure 3c-d**

| Gene name | Cortex Mean | Cortex SD | HC Mean | HC SD | p-value | Adj. p-value | Effect Size |
|---|---|---|---|---|---|---|---|
| ***Cacnb4*** | 0.790 | 1.621 | 0.000 | 0.000 | 0.000 | 0.021 | 0.558 |
| ***Cacng2*** | 0.226 | 0.584 | 0.000 | 0.000 | 0.003 | 0.246 | 0.442 |
| ***Kcnh3*** | 0.161 | 0.451 | 0.000 | 0.000 | 0.007 | 0.455 | 0.410 |
| ***Cacna1i*** | 0.177 | 0.497 | 0.000 | 0.000 | 0.007 | 0.455 | 0.409 |
| ***Kcnj4*** | 0.194 | 0.568 | 0.000 | 0.000 | 0.009 | 0.637 | 0.390 |
| ***Kcnc1*** | 0.290 | 0.687 | 0.050 | 0.224 | 0.019 | 1.000 | 0.394 |
| ***Cacna2d1*** | 1.145 | 1.587 | 0.550 | 0.686 | 0.022 | 1.000 | 0.417 |
| ***Cacng3*** | 0.081 | 0.275 | 0.000 | 0.000 | 0.024 | 1.000 | 0.336 |
| ***Kcnc4*** | 0.145 | 0.507 | 0.000 | 0.000 | 0.028 | 1.000 | 0.328 |
| ***Kcnh7*** | 0.210 | 0.750 | 0.000 | 0.000 | 0.031 | 1.000 | 0.320 |
| ***Cacna1a*** | 1.000 | 1.708 | 0.400 | 0.754 | 0.032 | 1.000 | 0.391 |
| ***Kcnb1*** | 1.274 | 2.847 | 0.500 | 0.761 | 0.056 | 1.000 | 0.308 |
| ***Kcnj2*** | 0.081 | 0.329 | 0.000 | 0.000 | 0.058 | 1.000 | 0.281 |
| ***Kcnj10*** | 0.387 | 1.107 | 0.100 | 0.308 | 0.070 | 1.000 | 0.294 |
| ***Kcnd3*** | 0.048 | 0.216 | 0.000 | 0.000 | 0.083 | 1.000 | 0.256 |
| ***Kcnn1*** | 0.081 | 0.375 | 0.000 | 0.000 | 0.096 | 1.000 | 0.246 |
| ***Cacna1e*** | 0.726 | 1.935 | 1.650 | 2.368 | 0.125 | 1.000 | 0.452 |
| ***Kcnk7*** | 0.081 | 0.417 | 0.000 | 0.000 | 0.133 | 1.000 | 0.222 |
| ***Kcnt1*** | 0.387 | 0.930 | 0.150 | 0.489 | 0.146 | 1.000 | 0.280 |
| ***Kcnf1*** | 0.065 | 0.356 | 0.000 | 0.000 | 0.159 | 1.000 | 0.207 |
| ***Kcnh4*** | 0.032 | 0.178 | 0.000 | 0.000 | 0.159 | 1.000 | 0.207 |
| ***Kcnq1*** | 0.032 | 0.178 | 0.000 | 0.000 | 0.159 | 1.000 | 0.207 |
| ***Kcnq5*** | 0.032 | 0.178 | 0.000 | 0.000 | 0.159 | 1.000 | 0.207 |
| ***Cacnb1*** | 0.032 | 0.178 | 0.000 | 0.000 | 0.159 | 1.000 | 0.207 |
| ***Kcnk13*** | 0.000 | 0.000 | 0.100 | 0.308 | 0.163 | 1.000 | 0.667 |
| ***Kcnj6*** | 0.065 | 0.248 | 0.300 | 0.733 | 0.173 | 1.000 | 0.564 |
| ***Cacna1h*** | 1.274 | 2.189 | 0.750 | 1.209 | 0.181 | 1.000 | 0.262 |
| ***Kcnq3*** | 0.048 | 0.282 | 0.000 | 0.000 | 0.182 | 1.000 | 0.196 |
| ***Kcna2*** | 0.500 | 0.901 | 0.300 | 0.470 | 0.203 | 1.000 | 0.244 |
| ***Kcnc3*** | 0.194 | 0.538 | 0.450 | 0.826 | 0.205 | 1.000 | 0.414 |
| ***Kcnh2*** | 0.065 | 0.400 | 0.000 | 0.000 | 0.208 | 1.000 | 0.185 |
| ***Kcnh5*** | 0.065 | 0.400 | 0.000 | 0.000 | 0.208 | 1.000 | 0.185 |
| ***Kcns2*** | 0.065 | 0.400 | 0.000 | 0.000 | 0.208 | 1.000 | 0.185 |
| ***Kcnj8*** | 0.210 | 0.656 | 0.650 | 1.496 | 0.215 | 1.000 | 0.475 |
| ***Kcnb2*** | 0.016 | 0.127 | 0.100 | 0.308 | 0.249 | 1.000 | 0.450 |
| ***Kcnc2*** | 0.016 | 0.127 | 0.100 | 0.308 | 0.249 | 1.000 | 0.450 |
| ***Kcnd2*** | 0.210 | 0.604 | 0.100 | 0.308 | 0.291 | 1.000 | 0.200 |
| ***Kcnma1*** | 0.710 | 1.419 | 0.400 | 1.095 | 0.314 | 1.000 | 0.230 |

| | | | | | | | |
|---|---|---|---|---|---|---|---|
| ***Kcnn4*** | 0.016 | 0.127 | 0.000 | 0.000 | 0.321 | 1.000 | 0.145 |
| ***Kcnu1*** | 0.032 | 0.254 | 0.000 | 0.000 | 0.321 | 1.000 | 0.145 |
| ***Kcnj13*** | 0.016 | 0.127 | 0.000 | 0.000 | 0.321 | 1.000 | 0.145 |
| ***Kcnj16*** | 0.016 | 0.127 | 0.000 | 0.000 | 0.321 | 1.000 | 0.145 |
| ***Kcnk6*** | 0.016 | 0.127 | 0.000 | 0.000 | 0.321 | 1.000 | 0.145 |
| ***Kcnk9*** | 0.016 | 0.127 | 0.000 | 0.000 | 0.321 | 1.000 | 0.145 |
| ***Kcnq4*** | 0.016 | 0.127 | 0.000 | 0.000 | 0.321 | 1.000 | 0.145 |
| ***Kcns3*** | 0.032 | 0.254 | 0.000 | 0.000 | 0.321 | 1.000 | 0.145 |
| ***Cacng5*** | 0.016 | 0.127 | 0.000 | 0.000 | 0.321 | 1.000 | 0.145 |
| ***Kcnj11*** | 0.000 | 0.000 | 0.050 | 0.224 | 0.330 | 1.000 | 0.459 |
| ***Kcna7*** | 0.000 | 0.000 | 0.050 | 0.224 | 0.330 | 1.000 | 0.459 |
| ***Cacng7*** | 0.000 | 0.000 | 0.050 | 0.224 | 0.330 | 1.000 | 0.459 |
| ***Cacng8*** | 0.000 | 0.000 | 0.050 | 0.224 | 0.330 | 1.000 | 0.459 |
| ***Cacna1f*** | 0.000 | 0.000 | 0.050 | 0.224 | 0.330 | 1.000 | 0.459 |
| ***Cacna2d2*** | 0.000 | 0.000 | 0.100 | 0.447 | 0.330 | 1.000 | 0.459 |
| ***Kcna1*** | 0.516 | 1.067 | 0.300 | 0.923 | 0.387 | 1.000 | 0.209 |
| ***Cacnb3*** | 0.323 | 0.901 | 0.200 | 0.410 | 0.406 | 1.000 | 0.151 |
| ***Cacnb2*** | 0.065 | 0.307 | 0.200 | 0.696 | 0.408 | 1.000 | 0.313 |
| ***Kcnv1*** | 0.113 | 0.483 | 0.050 | 0.224 | 0.429 | 1.000 | 0.145 |
| ***Kcnq2*** | 0.339 | 0.788 | 0.200 | 0.696 | 0.458 | 1.000 | 0.181 |
| ***Cacna1d*** | 0.710 | 1.475 | 0.500 | 0.946 | 0.461 | 1.000 | 0.153 |
| ***Cacna2d3*** | 0.194 | 0.698 | 0.100 | 0.447 | 0.487 | 1.000 | 0.145 |
| ***Kcna6*** | 0.032 | 0.178 | 0.100 | 0.447 | 0.516 | 1.000 | 0.253 |
| ***Cacna1c*** | 0.194 | 0.596 | 0.300 | 0.657 | 0.524 | 1.000 | 0.174 |
| ***Kcna5*** | 0.032 | 0.254 | 0.100 | 0.447 | 0.525 | 1.000 | 0.218 |
| ***Kcnk1*** | 0.081 | 0.329 | 0.050 | 0.224 | 0.640 | 1.000 | 0.100 |
| ***Kcnj12*** | 0.210 | 0.517 | 0.150 | 0.489 | 0.643 | 1.000 | 0.117 |
| ***Kcnt2*** | 0.129 | 0.383 | 0.100 | 0.308 | 0.732 | 1.000 | 0.079 |
| ***Cacna1g*** | 0.194 | 0.507 | 0.150 | 0.489 | 0.734 | 1.000 | 0.087 |
| ***Kcna4*** | 0.032 | 0.178 | 0.050 | 0.224 | 0.749 | 1.000 | 0.093 |
| ***Cacna1b*** | 0.032 | 0.254 | 0.050 | 0.224 | 0.767 | 1.000 | 0.072 |
| ***Kcnk3*** | 0.177 | 0.497 | 0.200 | 0.410 | 0.840 | 1.000 | 0.047 |
| ***Kcnn2*** | 0.113 | 0.367 | 0.100 | 0.308 | 0.877 | 1.000 | 0.036 |
| ***Kcnk2*** | 0.097 | 0.469 | 0.100 | 0.308 | 0.972 | 1.000 | 0.007 |

**c. Neurovascular signalling pathways:**

| Gene name | Cortex Mean | Cortex SD | HC Mean | HC SD | p-value | Adj. p-value | Effect Size |
|---|---|---|---|---|---|---|---|
| ***Cyp2j9*** | 0.226 | 0.838 | 0.000 | 0.000 | 0.038 | 0.683 | 0.309 |
| ***Grin2b*** | 1.871 | 3.123 | 0.800 | 1.609 | 0.050 | 0.846 | 0.377 |
| ***Nos1*** | 0.000 | 0.000 | 0.100 | 0.308 | 0.163 | 1.000 | 0.667 |
| ***Grin2c*** | 0.452 | 2.400 | 0.100 | 0.447 | 0.277 | 1.000 | 0.167 |
| ***Gucy1a2*** | 0.823 | 1.124 | 0.600 | 0.681 | 0.291 | 1.000 | 0.215 |
| ***Grin2d*** | 0.016 | 0.127 | 0.000 | 0.000 | 0.321 | 1.000 | 0.145 |
| ***Cyp2c50*** | 0.016 | 0.127 | 0.000 | 0.000 | 0.321 | 1.000 | 0.145 |
| ***Grin1*** | 0.661 | 1.414 | 0.450 | 0.826 | 0.415 | 1.000 | 0.163 |
| ***Grin2a*** | 0.065 | 0.248 | 0.150 | 0.489 | 0.461 | 1.000 | 0.266 |
| ***Gucy1b3*** | 1.452 | 2.434 | 1.800 | 1.908 | 0.512 | 1.000 | 0.150 |
| ***Gucy2g*** | 0.016 | 0.127 | 0.050 | 0.224 | 0.525 | 1.000 | 0.218 |

| | | | | | | | |
|---|---|---|---|---|---|---|---|
| ***Nos3*** | 0.435 | 1.350 | 0.300 | 0.733 | 0.570 | 1.000 | 0.110 |
| ***Cyp2j6*** | 0.129 | 0.424 | 0.100 | 0.308 | 0.741 | 1.000 | 0.073 |
| ***Ptges3l*** | 0.355 | 1.010 | 0.300 | 0.733 | 0.793 | 1.000 | 0.058 |
| ***Ptges*** | 0.065 | 0.400 | 0.050 | 0.224 | 0.839 | 1.000 | 0.040 |
| ***Gucy1a3*** | 1.597 | 2.336 | 1.500 | 2.090 | 0.862 | 1.000 | 0.042 |
| ***Ptges3*** | 0.629 | 0.962 | 0.650 | 1.040 | 0.937 | 1.000 | 0.021 |
| ***Ptges2*** | 0.048 | 0.216 | 0.050 | 0.224 | 0.978 | 1.000 | 0.007 |

#### Supplementary Table 5: Single cell mRNA comparisons for endothelial cells

a. **Ion channels:**

**See Figure 6h**

| Gene name | Cortex Mean | Cortex SD | HC Mean | HC SD | p-value | Adj. p-value | Effect Size |
|---|---|---|---|---|---|---|---|
| *Kcnj10* | 0.200 | 0.508 | 0.000 | 0.000 | 0.000 | 0.002 | 0.408 |
| *Kcnj2* | 0.184 | 0.530 | 0.000 | 0.000 | 0.000 | 0.012 | 0.360 |
| *Kcna1* | 0.376 | 1.148 | 0.000 | 0.000 | 0.000 | 0.027 | 0.339 |
| *Kcnh3* | 0.104 | 0.377 | 0.000 | 0.000 | 0.003 | 0.185 | 0.286 |
| *Cacnb3* | 0.080 | 0.301 | 0.000 | 0.000 | 0.004 | 0.253 | 0.276 |
| *Kcnk1* | 0.104 | 0.398 | 0.000 | 0.000 | 0.004 | 0.294 | 0.271 |
| *Kcnk3* | 0.056 | 0.231 | 0.000 | 0.000 | 0.008 | 0.534 | 0.251 |
| *Cacna1c* | 0.216 | 0.903 | 0.000 | 0.000 | 0.009 | 0.588 | 0.248 |
| *Cacng3* | 0.064 | 0.277 | 0.000 | 0.000 | 0.011 | 0.737 | 0.240 |
| *Cacnb4* | 0.720 | 2.684 | 0.100 | 0.316 | 0.019 | 1.000 | 0.239 |
| *Kcnj8* | 0.552 | 2.653 | 0.000 | 0.000 | 0.022 | 1.000 | 0.215 |
| *Kcnj12* | 0.040 | 0.197 | 0.000 | 0.000 | 0.025 | 1.000 | 0.211 |
| *Kcnh5* | 0.072 | 0.363 | 0.000 | 0.000 | 0.028 | 1.000 | 0.205 |
| *Kcna6* | 0.080 | 0.433 | 0.000 | 0.000 | 0.041 | 1.000 | 0.192 |
| *Kcnv1* | 0.080 | 0.433 | 0.000 | 0.000 | 0.041 | 1.000 | 0.192 |
| *Kcnma1* | 0.824 | 2.229 | 0.300 | 0.483 | 0.042 | 1.000 | 0.243 |
| *Kcnk7* | 0.032 | 0.177 | 0.000 | 0.000 | 0.045 | 1.000 | 0.188 |
| *Kcnc4* | 0.024 | 0.154 | 0.000 | 0.000 | 0.083 | 1.000 | 0.162 |
| *Kcnf1* | 0.048 | 0.307 | 0.000 | 0.000 | 0.083 | 1.000 | 0.162 |
| *Kcns2* | 0.024 | 0.154 | 0.000 | 0.000 | 0.083 | 1.000 | 0.162 |
| *Kcnn2* | 0.032 | 0.218 | 0.000 | 0.000 | 0.103 | 1.000 | 0.152 |
| *Kcnt2* | 0.032 | 0.218 | 0.000 | 0.000 | 0.103 | 1.000 | 0.152 |
| *Kcnj14* | 0.072 | 0.495 | 0.000 | 0.000 | 0.106 | 1.000 | 0.151 |
| *Kcnj4* | 0.144 | 0.998 | 0.000 | 0.000 | 0.109 | 1.000 | 0.149 |
| *Kcnj11* | 0.072 | 0.511 | 0.000 | 0.000 | 0.118 | 1.000 | 0.146 |
| *Cacna1i* | 0.040 | 0.295 | 0.000 | 0.000 | 0.132 | 1.000 | 0.140 |
| *Kcnn3* | 0.016 | 0.126 | 0.000 | 0.000 | 0.158 | 1.000 | 0.132 |
| *Kcnj16* | 0.016 | 0.126 | 0.000 | 0.000 | 0.158 | 1.000 | 0.132 |
| *Kcnj3* | 0.016 | 0.126 | 0.000 | 0.000 | 0.158 | 1.000 | 0.132 |
| *Kcnk10* | 0.016 | 0.126 | 0.000 | 0.000 | 0.158 | 1.000 | 0.132 |
| *Kcnk13* | 0.016 | 0.126 | 0.000 | 0.000 | 0.158 | 1.000 | 0.132 |
| *Kcnd1* | 0.016 | 0.126 | 0.000 | 0.000 | 0.158 | 1.000 | 0.132 |
| *Kcng2* | 0.016 | 0.126 | 0.000 | 0.000 | 0.158 | 1.000 | 0.132 |
| *Kcnh4* | 0.016 | 0.126 | 0.000 | 0.000 | 0.158 | 1.000 | 0.132 |
| *Kcnn1* | 0.024 | 0.199 | 0.000 | 0.000 | 0.181 | 1.000 | 0.125 |
| *Kcnu1* | 0.024 | 0.199 | 0.000 | 0.000 | 0.181 | 1.000 | 0.125 |
| *Cacna1e* | 0.552 | 1.316 | 2.500 | 4.275 | 0.184 | 1.000 | 1.153 |
| *Kcnc3* | 0.136 | 0.428 | 0.700 | 1.252 | 0.189 | 1.000 | 1.072 |
| *Kcnk2* | 0.128 | 0.457 | 0.400 | 0.699 | 0.255 | 1.000 | 0.569 |
| *Kcnd2* | 0.176 | 0.597 | 0.900 | 1.912 | 0.263 | 1.000 | 0.951 |
| *Cacng2* | 0.240 | 0.846 | 0.100 | 0.316 | 0.276 | 1.000 | 0.170 |
| *Kcnj15* | 0.016 | 0.179 | 0.000 | 0.000 | 0.319 | 1.000 | 0.093 |
| *Kcnj1* | 0.008 | 0.089 | 0.000 | 0.000 | 0.319 | 1.000 | 0.093 |

| | | | | | | | |
|---|---|---|---|---|---|---|---|
| ***Kcnk5*** | 0.016 | 0.179 | 0.000 | 0.000 | 0.319 | 1.000 | 0.093 |
| ***Kcna5*** | 0.016 | 0.179 | 0.000 | 0.000 | 0.319 | 1.000 | 0.093 |
| ***Kcng1*** | 0.008 | 0.089 | 0.000 | 0.000 | 0.319 | 1.000 | 0.093 |
| ***Kcng1.1*** | 0.008 | 0.089 | 0.000 | 0.000 | 0.319 | 1.000 | 0.093 |
| ***Kcnq4*** | 0.016 | 0.179 | 0.000 | 0.000 | 0.319 | 1.000 | 0.093 |
| ***Kcns1*** | 0.016 | 0.179 | 0.000 | 0.000 | 0.319 | 1.000 | 0.093 |
| ***Cacng4*** | 0.008 | 0.089 | 0.000 | 0.000 | 0.319 | 1.000 | 0.093 |
| ***Cacng5*** | 0.040 | 0.447 | 0.000 | 0.000 | 0.319 | 1.000 | 0.093 |
| ***Cacng8*** | 0.008 | 0.089 | 0.000 | 0.000 | 0.319 | 1.000 | 0.093 |
| ***Cacna2d2*** | 0.008 | 0.089 | 0.000 | 0.000 | 0.319 | 1.000 | 0.093 |
| ***Cacna1b*** | 0.008 | 0.089 | 0.100 | 0.316 | 0.383 | 1.000 | 0.771 |
| ***Cacna1d*** | 0.584 | 1.199 | 1.200 | 2.150 | 0.393 | 1.000 | 0.479 |
| ***Cacna1h*** | 0.208 | 0.873 | 0.100 | 0.316 | 0.404 | 1.000 | 0.127 |
| ***Cacna1a*** | 0.664 | 1.047 | 1.100 | 1.595 | 0.416 | 1.000 | 0.399 |
| ***Kcnb2*** | 0.016 | 0.126 | 0.100 | 0.316 | 0.425 | 1.000 | 0.572 |
| ***Kcna4*** | 0.024 | 0.154 | 0.100 | 0.316 | 0.470 | 1.000 | 0.448 |
| ***Kcnd3*** | 0.024 | 0.199 | 0.100 | 0.316 | 0.472 | 1.000 | 0.363 |
| ***Kcnq2*** | 0.240 | 0.865 | 0.500 | 1.080 | 0.475 | 1.000 | 0.295 |
| ***Kcnc1*** | 0.184 | 0.640 | 0.100 | 0.316 | 0.477 | 1.000 | 0.135 |
| ***Cacna2d3*** | 0.040 | 0.295 | 0.100 | 0.316 | 0.574 | 1.000 | 0.202 |
| ***Cacna2d1*** | 0.632 | 1.692 | 1.000 | 2.000 | 0.584 | 1.000 | 0.215 |
| ***Cacna1g*** | 0.120 | 0.548 | 0.200 | 0.422 | 0.584 | 1.000 | 0.148 |
| ***Kcnq3*** | 0.048 | 0.333 | 0.100 | 0.316 | 0.628 | 1.000 | 0.157 |
| ***Kcnj6*** | 0.144 | 0.564 | 0.100 | 0.316 | 0.700 | 1.000 | 0.080 |
| ***Kcnj9*** | 0.288 | 0.914 | 0.400 | 0.966 | 0.730 | 1.000 | 0.122 |
| ***Kcnt1*** | 0.128 | 0.508 | 0.200 | 0.632 | 0.733 | 1.000 | 0.139 |
| ***Kcnq1*** | 0.136 | 0.446 | 0.100 | 0.316 | 0.744 | 1.000 | 0.082 |
| ***Kcnc2*** | 0.136 | 0.497 | 0.100 | 0.316 | 0.748 | 1.000 | 0.074 |
| ***Kcnh7*** | 0.128 | 0.457 | 0.100 | 0.316 | 0.800 | 1.000 | 0.062 |
| ***Kcna2*** | 0.432 | 1.180 | 0.400 | 0.516 | 0.871 | 1.000 | 0.028 |
| ***Kcnb1*** | 0.768 | 2.342 | 0.700 | 1.252 | 0.881 | 1.000 | 0.030 |
| ***Kcnq5*** | 0.112 | 0.571 | 0.100 | 0.316 | 0.916 | 1.000 | 0.022 |
| ***Cacnb1*** | 0.104 | 0.521 | 0.100 | 0.316 | 0.972 | 1.000 | 0.008 |

**b. Neurovascular signalling pathways:**

**See Figure 6i, Supplementary Figures 3e-f**

| Gene name | Cortex Mean | Cortex SD | HC Mean | HC SD | p-value | Adj. p-value | Effect Size |
|---|---|---|---|---|---|---|---|
| ***Gucy1a2*** | 0.240 | 0.677 | 0.000 | 0.000 | 0.000 | 0.002 | 0.367 |
| ***Grin2c*** | 0.560 | 1.701 | 0.000 | 0.000 | 0.000 | 0.005 | 0.341 |
| ***Ptges*** | 0.176 | 0.540 | 0.000 | 0.000 | 0.000 | 0.005 | 0.338 |
| ***Cyp2j6*** | 0.168 | 0.632 | 0.000 | 0.000 | 0.004 | 0.046 | 0.275 |
| ***Ptges2*** | 0.056 | 0.231 | 0.000 | 0.000 | 0.008 | 0.092 | 0.251 |
| ***Gucy1a3*** | 0.264 | 1.212 | 0.000 | 0.000 | 0.016 | 0.180 | 0.226 |
| ***Gucy1b3*** | 0.408 | 1.232 | 0.100 | 0.316 | 0.045 | 0.450 | 0.258 |
| ***Ptges3l*** | 0.088 | 0.492 | 0.000 | 0.000 | 0.048 | 0.450 | 0.185 |
| ***Nos1*** | 0.016 | 0.126 | 0.000 | 0.000 | 0.158 | 1.000 | 0.132 |
| ***Grin2a*** | 0.048 | 0.307 | 0.500 | 0.972 | 0.176 | 1.000 | 1.159 |

| | | | | | | | |
|---|---|---|---|---|---|---|---|
| ***Ptges3*** | 0.432 | 0.892 | 0.900 | 0.994 | 0.179 | 1.000 | 0.520 |
| ***Nos3*** | 0.768 | 1.130 | 0.500 | 0.707 | 0.295 | 1.000 | 0.242 |
| ***Cyp2j5*** | 0.008 | 0.089 | 0.000 | 0.000 | 0.319 | 1.000 | 0.093 |
| ***Cyp2j9*** | 0.008 | 0.089 | 0.000 | 0.000 | 0.319 | 1.000 | 0.093 |
| ***Grin2b*** | 1.232 | 3.290 | 2.900 | 5.259 | 0.348 | 1.000 | 0.482 |
| ***Grin1*** | 0.456 | 1.428 | 0.900 | 1.912 | 0.489 | 1.000 | 0.303 |

#### Supplementary Table 6: Comparing average number of mRNA transcripts across the vascular beds of HC and neocortex

| Data from Figure/ Table | Mean | Standard Deviation | Test | Test Statistic | 95% Confidence Interval | Degrees of Freedom | P value |
|---|---|---|---|---|---|---|---|
| **Supplementary Data Table 4a: Mural Cells, Contractile Machinery** | HC: 2.0750 Ctx: 2.5692 | HC: 5.0452 Ctx: 7.0453 | Two-tailed unpaired t-test, unequal variances | t=0.3951 | -1.9925 to 2.9810 | df=85.17 | p=0.6937 |
| **Supplementary Data Table 4b: Mural Cells, Ion Channels** | HC: 0.1389 Ctx: 0.1956 | HC: 0.2483 Ctx: 0.2933 | Two-tailed unpaired t-test, unequal variances | t=1.2514 | -0.0329 to 0.1462 | df=138.23 | p=0.2129 |
| **Supplementary Data Table 4c: Mural Cells NV signalling** | HC: 0.3889 Ctx: 0.4919 | HC: 0.5220 Ctx: 0.5896 | Two-tailed unpaired t-test, unequal variances | t=0.5552 | -0.2744 to 0.4805 | df=33.5085 | p=0.5825 |
| **Supplementary Data Table 5a: Endothelial Cells, Ion Channels** | HC: 0.1592 Ctx: 0.1355 | HC: 0.3799 Ctx: 0.1893 | Two-tailed unpaired t-test, unequal variances | t=-0.4875 | -0.1202 to 0.0728 | df=110.09 | p=0.6269 |
| **Supplementary Data Table 5b: Endothelial Cells, NV signalling** | HC: 0.3625 Ctx: 0.3080 | HC: 0.7500 Ctx: 0.3331 | Two-tailed unpaired t-test, unequal variances | t=-0.2656 | -0.4815 to 0.3725 | df=20.6966 | p=0.7931 |
