## Supplementary material for "Hippocampus has lower oxygenation and weaker control of brain blood flow than cortex, due to microvascular differences": Statistics Reports

**Statistics Reports (SR1-8)**

The following tables report the mean, standard deviation, statistical test, test statistics, degrees of freedom and p-value for each statistical test reported in the main manuscript or supplementary figures, where not already reported in Supplementary Tables 1-6.

**SR1: Figure 2: Baseline haemodynamics in HC and V1 (comparison of response sizes)**

| Figure Label | Mean | Standard Deviation | Test | Test Statistic | 95% Confidence Interval | Degrees of Freedom | P value |
|---|---|---|---|---|---|---|---|
| **2a ($CMRO_2$)** | HC: 204.1<br>V1: 202.6 | HC: 58.5209<br>V1: 68.6911 | Two-tailed unpaired t-test | t=0.04816 | -65.22 to 62.32 | 16 | 0.9622 |
| **2b (CBF)** | HC: 298.8<br>V1: 392.1 | HC: 91.5472<br>V1: 47.3258 | Two-tailed unpaired t-test | t=2.551 | 15.34 to 171.2 | 15 | 0.0222* |
| **2c ($SO_2$)** | HC: 31.25<br>V1: 53.55 | HC: 3.4174<br>V1: 9.2856 | Two-tailed unpaired t-test | t=6.762 | 15.31 to 29.29 | 16 | 4.6E-6* |
| **2e (cap. density)** | HC: 1494<br>V1: 2802 | HC: 291.076<br>V1: 716.890 | Two-tailed unpaired t-test | t=4.116 | 589.1 to 2027 | 9 | 0.003* |
| **2g (diameter)** | HC: 5.490<br>V1: 5.305 | HC: 1.1425<br>V1: 1.0169 | Two-tailed unpaired t-test | t=0.7546 | -0.6726 to 0.3030 | 76 | 0.4528 |
| **2h (RBCV)** | HC: 1.210<br>V1: 2.592 | HC: 1.0551<br>V1: 2.2256 | Two-tailed unpaired t-test | t=3.505 | 0.5970 to 2.168 | 76 | 0.0008* |

**SR2a: Figure 3: Vessel responses to local neuronal calcium events (comparison of response sizes)**

| Figure Label | Mean | Standard Deviation | Test | Test Statistic | 95% Confidence Interval | Degrees of Freedom | P value |
|---|---|---|---|---|---|---|---|
| **3g (calcium)** | HC: 0.810<br>V1: 0.939 | HC: 1.7841<br>V1: 2.8760 | Two-tailed unpaired t-test | t=1.046 | -0.1051 to 0.3453 | 2015 | 0.2958 |
| **3h (diameter)** | HC: 0.019<br>V1: 0.025 | HC: 0.0222<br>V1: 0.0300 | Two-tailed unpaired t-test | t=4.863 | 0.00362 to 0.00851 | 2012 | 1.24E-6* |
| **3i (diameter, shuffled)** | HC: 0.004<br>V1: 0.004 | HC: 0.0040<br>V1: 0.0033 | Two-tailed unpaired t-test | t=0.5236 | -0.0002 to 0.00041 | 2015 | 0.6006 |
| **3l (calcium)** | HC: 0.829<br>V1: 0.721 | HC: 1.0428<br>V1: 1.0151 | Two-tailed unpaired t-test | t=0.9893 | -0.3245 to 0.1072 | 431 | 0.3231 |
| **3m (diameter)** | HC: 0.033<br>V1: 0.049 | HC: 0.0359<br>V1: 0.0424 | Two-tailed unpaired t-test | t=3.625 | 0.007253 to 0.02443 | 431 | 3.24E-4* |
| **3n ($NVC_{idx}$)** | HC: 0.094<br>V1: 0.201 | HC: 0.1492<br>V1: 0.2281 | Two-tailed unpaired t-test | t=4.785 | 0.06338 to 0.1517 | 431 | 2.35E-6* |

**SR2b: Figure 3: Vessel responses to local neuronal calcium events (comparison of response frequencies)**

| Figure Label | Test | Responsive | Non-Responsive | X-squared | Degrees of Freedom | P value |
|---|---|---|---|---|---|---|
| **3c (vessel responses)** | Pearson's Chi-squared test, with post-hoc pairwise comparisons | HC real: 120, V1 real: 313 | HC real: 659, V1 real: 925 | 27.092 | 1 | <0.0001* |

**Figure 4: Vasodilatory second messenger pathways in HC and cortex**
See Supplementary Data Tables 1-3.

**SR3: Figure 5: Wide-field neuronal activity patterns**

| Figure Label | Mean | Standard Deviation | Test | Test Statistic | 95% Confidence Interval | Degrees of Freedom | P value |
|---|---|---|---|---|---|---|---|
| **5c (correlation)** | HC: 0.017 V1: 0.0336 | HC: 0.0168 V1: 0.0319 | Two-tailed unpaired t-test | t=1.529 | -0.005679 to 0.03888 | 27 | 0.1379 |
| **5d (peak size)** | HC: 0.528 V1: 0.6598 | HC: 0.3155 V1: 0.2142 | Two-tailed unpaired t-test | t=1.338 | -0.07043 to 0.3344 | 27 | 0.1921 |
| **5h ($CMRO_2$)** | HC: 8.058 V1: 7.626 | HC: 13.1863 V1: 4.5496 | Two-tailed unpaired t-test | t=0.9719 | -1.302 to 0.4390 | 1920 | 0.3312 |
| **5j (Hbt)** | HC: 2.473 V1: 4.492 | HC: 1.4633 V1: 3.4391 | Two-tailed unpaired t-test | t=16.53 | 1.780 to 2.259 | 1920 | 1.8E-57* |

**SR4a: Figure 6: Pericyte morphology across brain regions (comparison of vessel sizes and ISDs)**

| Figure Label | Mean | Standard Deviation | Test | Test Statistic | 95% Confidence Interval | Degrees of Freedom | P value |
|---|---|---|---|---|---|---|---|
| **6a (diameters)** | HC: 2.76 V1: 2.99 | HC: 1.1599 V1: 1.4624 | Two-tailed unpaired t-test | t=1.236 | -0.1367 to 0.5969 | 214 | 0.2176 |
| **6b (pericyte ISD)** | HC: 102.5 V1: 86.21 | HC: 62.1564 V1: 40.7925 | Two-tailed unpaired t-test | t=2.322 | -30.08 to -2.457 | 214 | 0.0212* |

**SR4b: Figure 6: Pericyte morphology across brain regions (comparison of frequencies of cell types)**

| Figure Label | Test | Counts | X-squared | Degrees of Freedom | P value |
|---|---|---|---|---|---|
| **6d (mural cell type)** | Pearson's Chi-squared test | HC: SMC 18, EP 42, MP 172, TSP 98 V1: SMC 14, EP 61, MP 206, TSP 154 | 5.1936 | 7 | 0.158 |

**SR4c: Figure 6: Pericyte morphology across brain regions (comparison of cell lengths and diameters across regions)**

| Figure Label | Mean | Standard Deviation | Test | Test Statistic | Type III Sum of Squares | Degrees of Freedom | P value |
|---|---|---|---|---|---|---|---|
| **6e (diameter)** | HC: 3.85 V1: 3.81 | HC: 2.61 V1: 2.21 | Multifactorial ANOVA to compare region, cell type and region * cell type interaction | Region: F=1.271, Cell Type: F=393.887, Interaction: F=1.362 | Region: 2.815, Cell Type: 2616.616, Interaction: 9.049 | Region: df=1, Cell Type: df=3, Interaction: df=3 | Region: p=0.26, Cell Type: p=4.5E-154*, Interaction: p=0.253 |
| **6f (cell length)** | HC: 98.88 V1: 84.40 | HC: 46.66 V1: 35.10 | Multifactorial ANOVA to compare region, cell type and region * cell type interaction | Region: F=8.417, Cell Type: F=128.781, Interaction: F=4.102 | Region: 9077.352, Cell Type: 416672.2, Interaction: 13271.6 | Region: df=1, Cell Type: df=3, Interaction: df=3 | Region: p=0.004*, Cell Type: p=2.1E-67*, Interaction: p=0.007* |

The distribution of vessel diameters (p=0.02) and cell lengths (p<0.001) were unequal across brain regions (Mann-Whitney U test for independent samples), and the variance differed across groups. Therefore, we also ran multiple one-way ANOVA tests using Welch's post-hoc comparison, which also demonstrated significant effects of cell type on diameter and of region and cell type on cell length.

| Figure Label | Test | Test Statistic | P value |
|---|---|---|---|
| **6e (diameter)** | Welch statistic | Region: 0.04 Cell Type: 330.34 | Region: p=0.84 Cell Type: p=2.4E-57* |
| **6f (cell length)** | Welch statistic | Region: 22.23 Cell Type: 309.59 | Region: p=0.000003* Cell Type: p=3.38E-67* |

For Figures 6h-j, see Supplementary Data Tables 4-5.

**SR5a: Supplementary Figure 1: The contribution of cellular input to vascular responses (comparison of ROI type frequencies)**

| Figure Label | Test | Counts | X-squared | Degrees of Freedom | P value |
|---|---|---|---|---|---|
| **SD 1b (ROI type distribution)** | Pearson's Chi-squared test, 2x2 contingency table | HC >60% NP: 30 HC >60% Soma: 10 V1 >60% NP: 32 V1 >60% Soma: 9 | 0.105 | df=3 | p=0.7976 |

**SR5b: Supplementary Figure 1: The contribution of cellular input to vascular responses (comparison of response frequencies)**

| Figure Label | Test | Responsive | Non-Responsive | X-squared | Degrees of Freedom | P value |
|---|---|---|---|---|---|---|
| **SD 1d (vessel responses)** | Pearson's Chi-squared test, with post-hoc pairwise table | HC >60% NP: 66 HC >60% Soma: 39 V1 >60% NP: 271 | HC >60% NP: 401 HC >60% Soma: 175 V1 >60% NP: 737 V1 | 34.707 | df=3 | p=1.4E-7* |

| | | | |
|---|---|---|---|
| | | V1 >60% Soma: 42 | >60% Soma: 188 |

| Comparison | P value | Adjusted P Value |
|---|---|---|
| **HC >60% NP vs. HC >60% Soma** | 0.172 | 0.217 |
| **V1 >60% NP vs. V1 >60% Soma** | 0.007 | 0.0148* |
| **HC >60% Soma vs. V1 >60% Soma** | 1.00 | 1.00 |
| **HC >60% NP vs. V1 >60% NP** | 2.69E-8 | 1.61E-7* |

#### SR5c: Supplementary Figure 1: The contribution of cellular input to vascular responses (comparison of response sizes)

| Figure Label | Mean | Standard Deviation | Test | Test Statistic | 95% Confidence Interval | Degrees of Freedom | P value |
|---|---|---|---|---|---|---|---|
| **SD 1c (diameter)** | HC >60% NP: 8.8, HC >60% Soma: 10.6, V1 >60% NP: 7.9, V1 >60% Soma: 10 | HC >60% NP: 3.8, HC >60% Soma: 5, V1 >60% NP: 4, V1 >60% Soma: 3.9 | Two-tailed unpaired t-test | HC: t=1.182 V1: t=1.417 | HC: -1.263 to 4.804 V1: -0.9134 to 5.191 | HC: df=38 V1: df=39 | HC: p=0.2447 V1: p=0.1643 |
| **SD 1e (HC calcium)** | >60% NP: 0.69 >60% Soma: 0.95 | >60% NP: 0.72 >60% Soma: 1.50 | Two-tailed unpaired t-test | t=1.131 | -0.1921 to 0.7011 | df=97 | p=0.2608 |
| **SD 1f (HC diameter)** | >60% NP: 0.023 >60% Soma: 0.018 | >60% NP: 0.016 >60% Soma: 0.014 | Two-tailed unpaired t-test | t=1.860 | -0.01200 to 0.0003895 | df=97 | p=0.0659 |
| **SD 1g (HC $NVC_{idx}$)** | >60% NP: 0.090 >60% Soma: 0.041 | >60% NP: 0.179 >60% Soma: 0.031 | Two-tailed unpaired t-test | t=1.664 | -0.1060 to 0.009305 | df=97 | p=0.0993 |
| **SD 1h (V1 calcium)** | >60% NP: 0.735 >60% Soma: 0.481 | >60% NP: 1.066 >60% Soma: 0.512 | Two-tailed unpaired t-test | t=1.511 | -0.5831 to 0.07646 | df=311 | p=0.1317 |
| **SD 1i (V1 diameter)** | >60% NP: 0.042 >60% Soma: 0.049 | >60% NP: 0.039 >60% Soma: 0.060 | Two-tailed unpaired t-test | t=1.004 | -0.006716 to 0.02071 | df=311 | p=0.3162 |
| **SD 1j (V1 $NVC_{idx}$)** | >60% NP: 0.186 >60% Soma: 0.158 | >60% NP: 0.220 >60% Soma: 0.170 | Two-tailed unpaired t-test | t=0.7921 | -0.09787 to 0.04169 | df=311 | p=0.4289 |

#### SR5d: Supplementary Figure 1: The contribution of cellular input to vascular responses (comparison of response sizes across region and ROI type)

| Figure Label | Test | Test Statistic | Type III Sum of Squares | Degrees of Freedom | P value |
|---|---|---|---|---|---|
| **SD 1e, 1h (calcium)** | Multifactorial ANOVA: region, ROI type, region * ROI type interaction | Region: F=2.419, ROI Type: F=0.00002, Interaction: F=3.475 | Region: 2.572, ROI Type: 2E-5, Interaction: 3.695 | Region: df=1, ROI Type: df=1, Interaction: df=1 | Region: p=0.121, ROI Type: p=0.997, Interaction: p=0.063 |
| **SD 1f, 1i (diameter)** | Multifactorial ANOVA: region, ROI type, region | Region: F=26.374, ROI Type: F=0.015, | Region: 0.037, ROI Type: 2.03E-5, | Region: df=1, ROI Type: df=1, | Region: p=4.4E-7*, ROI Type: |

| | | | | | |
|---|---|---|---|---|---|
| | * ROI type interaction | Interaction: F=1.676 | Interaction: 0.002 | Interaction: df=1 | p=0.904, Interaction: p=0.196 |
| **SD 1g, 1j ($NVC_{idx}$)** | Multifactorial ANOVA: region, ROI type, region * ROI type interaction | Region: F=16.374, ROI Type: F=2.113, Interaction: F=0.148 | Region: 0.648, ROI Type: 0.084, Interaction: 0.006 | Region: df=1, ROI Type: df=1, Interaction: df=1 | Region: p=6E-4*, ROI Type: p=0.147 Interaction: p=0.700 |

**SR6a: Supplementary Figure 2: The contribution of laminar organisation to vascular responses (comparison of response frequencies)**

| Figure Label | Test | Responsive | Non-Responsive | X-squared | Degrees of Freedom | P value |
|---|---|---|---|---|---|---|
| **SD 2a (HC vessel responses)** | Pearson's Chi-squared test, with post-hoc pairwise table | SO: 12<br>SP: 76<br>SR: 24<br>SLM: 8 | SO: 80<br>SP: 391<br>SR: 133<br>SLM: 55 | 1.0202 | df=3 | p=0.7964 |

| Comparison | P value | Adjusted P Value |
|---|---|---|
| **SO vs. SP** | 0.535 | 1.000 |
| **SO vs. SR** | 0.765 | 1.000 |
| **SO vs. SLM** | 1.000 | 1.000 |
| **SP vs. SR** | 0.868 | 1.000 |
| **SP vs. SLM** | 0.585 | 1.000 |
| **SR vs. SLM** | 0.779 | 1.000 |

**SR6b: Supplementary Figure 2: The contribution of laminar organisation to vascular responses (comparison of frequency of observations in each layer)**

| Figure Label | Test | Responsive | Non-Responsive | X-squared | Degrees of Freedom | P value |
|---|---|---|---|---|---|---|
| **SD 2c (V1 vessel responses)** | Pearson's Chi-squared test, 2x2 contingency table | L1: 48<br>L2/3: 261 | L1: 164<br>L2/3: 747 | 0.81469 | df=1 | p=0.3667 |

**SR6c: Supplementary Figure 2: The contribution of laminar organisation to vascular responses (comparison of vessel sizes across HC layers)**

| Figure Label | Mean | Standard Deviation | Test | Test Statistic | Degrees of Freedom | P value |
|---|---|---|---|---|---|---|
| **SD 2b (HC vessel sizes)** | SO: 9.6914<br>SP: 8.6245<br>SR: 8.1862<br>SLM: 9.117 | SO: 3.4501<br>SP: 3.2296<br>SR: 3.7051<br>SLM: 5.4891 | One-way ANOVA with Bonferroni post-hoc comparisons | F=0.2487 | df=3 | p=0.8617 |

| Comparison | 95% Confidence Interval | Adjusted P Value |
|---|---|---|
| **SO vs. SP** | -3.276 to 5.410 | >0.999 |
| **SO vs. SR** | -3.675 to 6.685 | >0.999 |
| **SO vs. SLM** | -5.699 to 6.847 | >0.999 |
| **SP vs. SR** | -3.694 to 4.570 | >0.999 |
| **SP vs. SLM** | -5.933 to 4.947 | >0.999 |
| **SR vs. SLM** | -7.060 to 5.198 | >0.999 |

**SR6d: Supplementary Figure 2: The contribution of laminar organisation to vascular responses (comparison of calcium response sizes in HC)**

| Figure Label | Mean | Standard Deviation | Test | Test Statistic | Degrees of Freedom | P value |
|---|---|---|---|---|---|---|
| **SD 2e (HC calcium)** | SO: 1.1780<br>SP: 0.7840<br>SR: 0.6604<br>SLM: 0.5236 | SO: 1.0884<br>SP: 1.1693<br>SR: 0.5042<br>SLM: 0.4752 | One-way ANOVA with Bonferroni post-hoc comparisons | F=0.8710 | df=3 | p=0.5154 |

| Comparison | 95% Confidence Interval | Adjusted P Value |
|---|---|---|
| **SO vs. SP** | -0.4647 to 1.253 | >0.9999 |
| **SO vs. SR** | -0.4597 to 1.495 | 0.9471 |
| **SO vs. SLM** | -0.6072 to 1.916 | 0.9990 |
| **SP vs. SR** | -0.5236 to 0.7708 | >0.9999 |
| **SP vs. SLM** | -0.7669 to 1.288 | >0.9999 |
| **SR vs. SLM** | -0.9916 to 1.265 | >0.9999 |

**SR6e: Supplementary Figure 2: The contribution of laminar organisation to vascular responses (comparison of vascular response sizes in HC)**

| Figure Label | Mean | Standard Deviation | Test | Test Statistic | Degrees of Freedom | P value |
|---|---|---|---|---|---|---|
| **SD 2f (HC diameter)** | SO: 0.0586<br>SP: 0.0278<br>SR: 0.0503<br>SLM: 0.0454 | SO: 0.0693<br>SP: 0.0207<br>SR: 0.0445<br>SLM: 0.0368 | One-way ANOVA with Bonferroni post-hoc comparisons | F=4.630 | df=3 | p=0.0043* |

| Comparison | 95% Confidence Interval | Adjusted P Value |
|---|---|---|
| **SO vs. SP** | 0.001798 to 0.05974 | 0.0310* |
| **SO vs. SR** | -0.02466 to 0.04129 | >0.9999 |
| **SO vs. SLM** | -0.02937 to 0.05577 | >0.9999 |
| **SP vs. SR** | -0.04429 to -0.0006177 | 0.0403* |
| **SP vs. SLM** | -0.05223 to 0.01710 | >0.9999 |
| **SR vs. SLM** | -0.03319 to 0.04296 | >0.9999 |

**SR6g: Supplementary Figure 2: The contribution of laminar organisation to vascular responses (comparison of $NVC_{idx}$ in HC)**

| Figure Label | Mean | Standard Deviation | Test | Test Statistic | Degrees of Freedom | P value |
|---|---|---|---|---|---|---|
| **SD 2g (HC $NVC_{idx}$)** | SO: 0.0775<br>SP: 0.1149<br>SR: 0.1251<br>SLM: 0.1182 | SO: 0.0759<br>SP: 0.1827<br>SR: 0.1302<br>SLM: 0.1318 | One-way ANOVA with Bonferroni post-hoc comparisons | F=0.2403 | df=3 | p=0.8681 |

| Comparison | 95% Confidence Interval | Adjusted P Value |
|---|---|---|
| **SO vs. SP** | -0.1733 to 0.09839 | >0.9999 |
| **SO vs. SR** | -0.2022 to 0.1070 | >0.9999 |
| **SO vs. SLM** | -0.2403 to 0.1589 | >0.9999 |
| **SP vs. SR** | -0.1126 to 0.09219 | >0.9999 |
| **SP vs. SLM** | -0.1658 to 0.1592 | >0.9999 |
| **SR vs. SLM** | -0.1716 to 0.1854 | >0.9999 |

**SR6h: Supplementary Figure 2: The contribution of laminar organisation to vascular responses (comparison of response sizes in V1)**

| Figure Label | Mean | Standard Deviation | Test | Test Statistic | 95% Confidence Interval | Degrees of Freedom | P value |
|---|---|---|---|---|---|---|---|
| **SD 2d** | L1: 9.7921 | L1: 4.8976 | Two-tailed unpaired t-test | t=1.118 | -5.075 to 1.463 | df=38 | p=0.2704 |

| | | | | | | | |
|---|---|---|---|---|---|---|---|
| **(V1 vessel sizes)** | L2/3: 7.9863 | L2/3: 3.8780 | | | | | |
| **SD 2h (V1 calcium)** | L1: 0.6273 L2/3: 0.6876 | L1: 1.0405 L2/3: 0.9847 | Two-tailed unpaired t-test | t=0.3864 | -0.2467 to 0.3673 | df=307 | p=0.6995* |
| **SD 2i (V1 diameter)** | L1: 0.0778 L2/3: 0.0477 | L1: 0.0524 L2/3: 0.0396 | Two-tailed unpaired t-test | t=4.589 | -0.04303 to -0.0172 | df=307 | p=6.5E-6* |
| **SD 2j (V1 $NVC_{idx}$)** | L1: 0.4181 L2/3: 0.1907 | L1: 0.3335 L2/3: 0.2230 | Two-tailed unpaired t-test | t=5.956 | -0.3026 to -0.1523 | df=307 | p=7.1E-9* |

#### SR6i: Supplementary Figure 2: The contribution of laminar organisation to vascular responses (comparison of response sizes between regions)

| Figure Label | Test | Test Statistic | Sum of Squares | Degrees of Freedom | P value |
|---|---|---|---|---|---|
| **SD 2g, 2j ($NVC_{idx}$)** | One-way ANOVA with Bonferroni post-hoc comparisons | F=32.011 | 3.193 | 2 | p=1.12E-13 |

| Comparison | 95% Confidence Interval | P Value |
|---|---|---|
| **HC vs. L1** | -0.3964 to -0.2131 | 3.85E-14* |
| **HC vs. L2/3** | -0.1365 to -0.0181 | 0.005* |
| **L1 vs. L2/3** | 0.1431 to 0.3117 | 7.43E-10* |

#### Supplementary Figure 3: Neurovascular pathways in vascular cells

For Figures 3a-f, see Supplementary Data Tables 4-5.

#### SR7a: Supplementary Figure 4: The impact of hippocampal cranial window surgery (comparison of response sizes)

| Figure Label | Mean | Standard Deviation | Test | Test Statistic | 95% Confidence Interval | Degrees of Freedom | P value |
|---|---|---|---|---|---|---|---|
| **SD 4b (behaviour)** | HC: 65.8 V1: 67 | HC: 11.09 V1: 15.41 | Two-tailed unpaired t-test | t=-0.214 | -13.88 to 11.48 | 9 | 0.8352 |
| **SD 4e (vessel sizes)** | HC <100µm: 9.23, HC >110µm: 9.72 V1 <100µm: 9.08, V1 >110µm: 7.79 | HC <100µm: 3.87, HC >110µm: 4.61 V1 <100µm: 4.77, V1 >110µm: 3.46 | Two-tailed unpaired t-test | HC: t=0.3748 V1: t=1.003 | HC: -2.171 to 3.163 V1: -3.884 to 1.308 | HC: df=43 V1: df=39 | HC: 0.71 V1: 0.32 |
| **SD 4g (HC calcium)** | <100µm: 0.84 >110µm: 0.68 | <100µm: 1.21 >110µm: 0.52 | Two-tailed unpaired t-test | t=0.8136 | -0.5536 to 0.2312 | 118 | 0.42 |
| **SD 4h (HC diameter)** | <100µm: 0.03 >110µm: 0.04 | <100µm: 0.03 >110µm: 0.04 | Two-tailed unpaired t-test | t=1.534 | -0.003099 to 0.02442 | 118 | 0.13 |
| **SD 4i (HC $NVC_{idx}$)** | <100µm: 0.10 >110µm: 0.13 | <100µm: 0.12 >110µm: 0.22 | Two-tailed unpaired t-test | t=0.8759 | -0.03433 to 0.08878 | 118 | 0.38 |

| | | | | | | | |
|---|---|---|---|---|---|---|---|
| **SD 4j (V1 calcium)** | <100μm: 0.73 >110μm: 0.69 | <100μm: 1.02 >110μm: 1.01 | Two-tailed unpaired t-test | t=0.3440 | -0.2782 to 0.1954 | 311 | 0.73 |
| **SD 4k (V1 diameter)** | <100μm: 0.05 >110μm: 0.05 | <100μm: 0.05 >110μm: 0.04 | Two-tailed unpaired t-test | t=0.4181 | -0.01217 to 0.007906 | 311 | 0.68 |
| **SD 4l (V1 $NVC_{idx}$)** | <100μm: 0.24 >110μm: 0.22 | <100μm: 0.28 >110μm: 0.24 | Two-tailed unpaired t-test | t=0.6009 | -0.07805 to 0.04153 | 311 | 0.55 |

### SR7b: Supplementary Figure 4: The impact of hippocampal cranial window surgery (comparison of vessel depth distribution)

| **Figure Label** | **Test** | **Counts** | **X-squared** | **Degrees of Freedom** | **P value** |
|---|---|---|---|---|---|
| **SD 4d (depth distribution)** | Pearson's Chi-squared test | HC: <100μm 81, >110μm 36<br>V1: <100μm 130, >110μm 183 | 26.1423 | 3 | <0.00001* |

### SR7c: Supplementary Figure 4: The impact of hippocampal cranial window surgery (comparison of response frequencies)

| **Figure Label** | **Test** | **Responsive** | **Non-Responsive** | **X-squared** | **Degrees of Freedom** | **P value** |
|---|---|---|---|---|---|---|
| **SD 4f (vessel responses)** | Pearson's Chi-squared test, with post-hoc pairwise table | HC <100μm: 79, HC >110μm: 35, V1 <100μm: 109, V1 >110μm: 204 | HC <100μm: 448, HC >110μm: 186,<br>V1 <100μm: 363, V1 >110μm: 562 | 30.092 | 3 | 1.32E-6* |

| **Comparison** | **P value** | **Adjusted P Value** |
|---|---|---|
| **HC <100μm vs. HC >110μm** | 0.855 | 0.855 |
| **HC <100μm vs. V1 <100μm** | 0.00142 | 0.00284* |
| **HC >110μm vs. V1 >110μm** | 0.00132 | 0.00284* |
| **V1 <100μm vs. V1 >110μm** | 0.185 | 0.222 |

### SR8: Supplementary Figure 5: Testing the reliability of the Oxy-CBF probe (comparison of response sizes)

| **Figure Label** | **Mean** | **Standard Deviation** | **Test** | **Test Statistic** | **95% Confidence Interval** | **Degrees of Freedom** | **P value** |
|---|---|---|---|---|---|---|---|
| **SD 5b (resting $CMRO_2$, CBF and $SO_2$)** | $CMRO_2$ – WT: 190.6 FL: 180.59<br>CBF – WT: 442.5 FL: 400.34<br>$SO_2$ – WT: 56.08 FL: 54.48 | $CMRO_2$ – WT: 50.58 FL: 36.65<br>CBF – WT: 131.1 FL: 50.88<br>$SO_2$ – WT: 3.78 FL: 4.72 | Two-tailed unpaired t-test | $CMRO_2$: t=0.3954, CBF: t=0.8237, $SO_2$: t=0.5864 | $CMRO_2$: -66.40 to 46.38, CBF: -156.0 to 71.81 $SO_2$: -7.693 to 4.487 | df=10 | $CMRO_2$: p=0.7008, CBF: p=0.4293, $SO_2$: p=0.5706 |
| **SD 5c** | CBF – | CBF – WT: 2.89 FL: 5.61 | Two-tailed | CBF: t=0.5978 | CBF: -6.318 to 10.15 | df=5 | CBF: p=0.5760 |

| | | | | | | | |
|---|---|---|---|---|---|---|---|
| **($CMRO_2$, CBF and $SO_2$ peaks)** | WT: 10.72 FL: 12.63 $SO_2$ – WT: 6.66 FL: 6.39 Hbt – WT: 4.30 FL: 4.61 | $SO_2$ – WT: 1.48 FL: 1.90 Hbt – WT: 1.45 FL: 1.90 | unpaired t-test | $SO_2$: t=0.1046 Hbt: t=0.2438 | $SO_2$: -6.773 to 6.244 Hbt: -2.923 to 3.536 | | $SO_2$: p=0.9207 Hbt: P=0.8170 |
| **SD 5d ($CMRO_2$, CBF and $SO_2$ pre- and post-pentobarbital injection)** | CMRO2 – Pre: 266.9 Post: 5.65 CBF – Pre: 540.5 Post: 5.65 $SO_2$ – Pre: 51.49 Post: 0 | CMRO2 – Pre: 95.77 Post: 3.78 CBF – Pre: 149.4 Post: 3.78 $SO_2$ – Pre: 5.27 Post: 0 | Two-tailed unpaired t-test | CMRO2: t=4.720 CBF: t=6.200 SO2: t=16.93 | CMRO2: -414.9 to -107.6 CBF: -774.3 to -295.3 $SO_2$: -59.94 to -43.05 | df=4 | CMRO2: p=0.0092* CBF: p=0.0034* $SO_2$: p=7.1E-5* |
